# DELE1 promotes translation-associated homeostasis, growth, and survival in mitochondrial myopathy

**DOI:** 10.1101/2024.02.29.582673

**Authors:** Hsin-Pin Lin, Jennifer D. Petersen, Alexandra J. Gilsrud, Angelo Madruga, Theresa M. D’Silva, Xiaoping Huang, Mario K. Shammas, Nicholas P. Randolph, Yan Li, Drew R. Jones, Michael E. Pacold, Derek P. Narendra

## Abstract

Mitochondrial dysfunction causes devastating disorders, including mitochondrial myopathy. Here, we identified that diverse mitochondrial myopathy models elicit a protective mitochondrial integrated stress response (mt-ISR), mediated by OMA1-DELE1 signaling. The response was similar following disruptions in mtDNA maintenance, from knockout of *Tfam*, and mitochondrial protein unfolding, from disease-causing mutations in CHCHD10 (G58R and S59L). The preponderance of the response was directed at upregulating pathways for aminoacyl-tRNA biosynthesis, the intermediates for protein synthesis, and was similar in heart and skeletal muscle but more limited in brown adipose challenged with cold stress. Strikingly, models with early DELE1 mt-ISR activation failed to grow and survive to adulthood in the absence of *Dele1*, accounting for some but not all of OMA1’s protection. Notably, the DELE1 mt-ISR did not slow net protein synthesis in stressed striated muscle, but instead prevented loss of translation-associated proteostasis in muscle fibers. Together our findings identify that the DELE1 mt-ISR mediates a stereotyped response to diverse forms of mitochondrial stress and is particularly critical for maintaining growth and survival in early-onset mitochondrial myopathy.

## Introduction

Mitochondria generate energy through oxidative phosphorylation (OXPHOS) (Classics Mitchell, 1966). They are also important for cell growth and maintenance, compartmentalizing amino acid synthesis and one carbon (1C) metabolism pathways to generate biosynthetic intermediates (Zong *et al*, 2016). The biosynthetic and energetic functions of mitochondria are coupled through redox equivalents such as NADH, and cell growth can be limited by OXPHOS inhibition even in the setting of sufficient ATP production (Luengo *et al*, 2021).

Maintaining optimal OXPHOS is thus important for cell maintenance and growth but is also complicated by the intricacy of the OXPHOS system. In OXPHOS, energy from fuels like glucose and fatty acids are initially transferred to redox equivalents by the tricyclic acid (TCA) cycle (and other pathways) and fed into the electron transport chain (ETC), composed of four inner mitochondrial membrane (IMM) embedded complexes (Chandel, 2021). The ETC uses redox energy from these reducing equivalents to pump protons across the IMM. The IMM stores this potential energy in an electrochemical gradient, which then tunnels protons through the F_1_F_O_-ATP synthase to drive ATP synthesis. Damage to any component of this IMM-embedded system, including the IMM itself, can potentially hobble the mitochondrial production of ATP and biosynthetic intermediates.

Indeed, mutations in over 250 genes cause primary mitochondrial disorders, presenting often as mitochondrial myopathy (Mayr *et al*, 2015), and often with abnormal growth and short stature (Boal *et al*, 2019). Some of these likely damage the IMM through protein unfolding stress, as we and others recently demonstrated for mutations in the nuclear DNA (nDNA)-encoded intermembrane space (IMS) protein CHCHD10, causing Dominant Inherited Mitochondrial Myopathy (IMMD) (Shammas *et al*, 2022; Ajroud-Driss *et al*, 2015; Bannwarth *et al*, 2014; Genin *et al*, 2019; Anderson *et al*, 2019). Others cause severe OXPHOS deficiency through either mutations in the mitochondrial DNA (mtDNA) itself or the machinery needed to express the thirteen mtDNA-encoded OXPHOS subunits (Zeviani *et al*, 1989; Holt *et al*, 1988; Spelbrink *et al*, 2001; Goethem *et al*, 2001). Mutations affecting mtDNA or its expression account for most mitochondrial myopathy cases (Gorman *et al*, 2015). The mechanisms of mitochondrial damage can thus be diverse, but they likely have in common disruption of OXPHOS and its coupled biosynthetic pathways. Patient survival may depend on how well striated muscle adapts to these disruptions in mitochondrial metabolism (Hathazi *et al*, 2020).

One form of adaptation to mitochondrial stress is mediated through retrograde mitochondria-to-nucleus (mitonuclear) signaling. The first mitonuclear signal in mammalian cells was identified in the setting of mitochondrial protein unfolding stress and was named the mitochondrial unfolded protein response (mt-UPR) (Zhao *et al*, 2002). Subsequently, an amino acid starvation-like response was identified in mouse models with defects in mtDNA maintenance (e.g., mutations in *Twinkle* or *Tfam*) or mtDNA expression (e.g., mutations in *Dars2*) causing severe OXPHOS deficiency (Tyynismaa *et al*, 2010; Dogan *et al*, 2014; Kühl *et al*, 2017). Responses like these were initially attributed to energy and amino acid sensing pathways outside of the mitochondria, with upstream signaling from AKT, mTOR, and/or GCN2, converging on transcription factors such as ATF4, ATF5, and CHOP (Tyynismaa *et al*, 2010; Khan *et al*, 2017; Mick *et al*, 2020). Related work emphasized the imbalance between mtDNA and nDNA encoded OXPHOS subunits in the setting of diminished mtDNA expression as a mitonuclear signal (Houtkooper *et al*, 2013). However, it is not known whether these responses to different forms of mitochondrial stress are mediated by a common molecular signaling pathway *in vivo* or if they reflect distinct molecular pathways signaling distinct stresses, such as mitochondrial protein unfolding vs. mitonuclear imbalance.

In cultured cells, one of these mitonuclear signals is the OMA1-DELE1-HRI-eIF2α-ATF4 signaling cascade (hereafter, the DELE1 mt-ISR) (Fessler *et al*, 2020; Guo *et al*, 2020). Recently, we demonstrated that the OMA1-DELE1 pathway can also signal the mt-ISR *in vivo*, in response to protein misfolding of CHCHD10 with the IMMD-causing mutation (Shammas *et al*, 2022). We additionally found that the OMA1-dependent stress response is strongly protective in the IMMD model. Similar findings were independently reported for two other cardiomyopathy models (one from conditional knockout (KO) of a Complex IV subunit, COX10 (Ahola *et al*, 2022), and the other from KO of the cardiolipin remodeling protein Tafazzin (Huynh *et al*, 2022; Zhu *et al*, 2022)), together supporting the idea that the DELE1 mt-ISR mediates at least some mitochondria-to-nucleus signaling in response to mitochondrial stress *in vivo*. However, it is not known whether there is a common OMA1-DELE1 stress pathway responding to diverse mitochondrial stressors, such as protein unfolding and mtDNA maintenance. Additionally, the mt-ISR has not been examined in skeletal muscle, the tissue that is primarily affected in mitochondrial myopathies.

Here, we provide a comprehensive view of the OMA1-DELE1 stress response *in vivo*, using a novel *Dele1* KO mouse subjected to four diverse mitochondrial stressors in striated muscle, including protein unfolding stress (mutant CHCHD10 protein unfolding) and severe mitonuclear imbalance from a defect in mtDNA maintenance (*Tfam* KO) (Hansson *et al*, 2004). Using a multi-omics approach, we identified that DELE1 mediates a stereotyped transcriptional response to these diverse mitochondrial stressors to maintain biosynthetic pathways, particularly protein synthesis. This response was similar in cardiac and skeletal muscle and partially overlapped with the physiologic DELE1-dependent response we uncovered in brown adipose tissues (BAT) subjected to cold stress. Although protective in all myopathy/cardiomyopathy models tested, the DELE1 mt-ISR provided dramatically increased survival in two myopathy/cardiomyopathy models with stress onset during early postnatal growth, underscoring the importance of the mt-ISR for promoting anabolism in the face of mitochondrial stress. Importantly, the DELE1 mt-ISR signature was observed in skeletal muscle from patients with mitochondrial myopathy from several prior independent studies, together suggesting that the DELE1 mt-ISR is a critical response for maintaining striated muscle in mitochondrial myopathy.

## Results

### DELE1 mediates the mt-ISR in response to physiologic stress

We previously found that OMA1 promotes survival in a model of early-onset mitochondrial myopathy (Shammas *et al*, 2022). OMA1 activates the mt-ISR by cleaving DELE1, generating a short form of DELE1 (S-DELE1) (Fessler *et al*, 2020; Guo *et al*, 2020). S-DELE1, in turn, activates HRI to phosphorylate eIF2α, thereby triggering the ISR (Fig. 1A, top). The ISR acutely slows protein synthesis overall, by decreasing the rate of translation initiation, but allows for the preferential translation of transcription factors with upstream open reading frames such as ATF4, causing global changes in gene expression (Pakos-Zebrucka *et al*, 2016; Wek, 2018).

**Figure 1.**
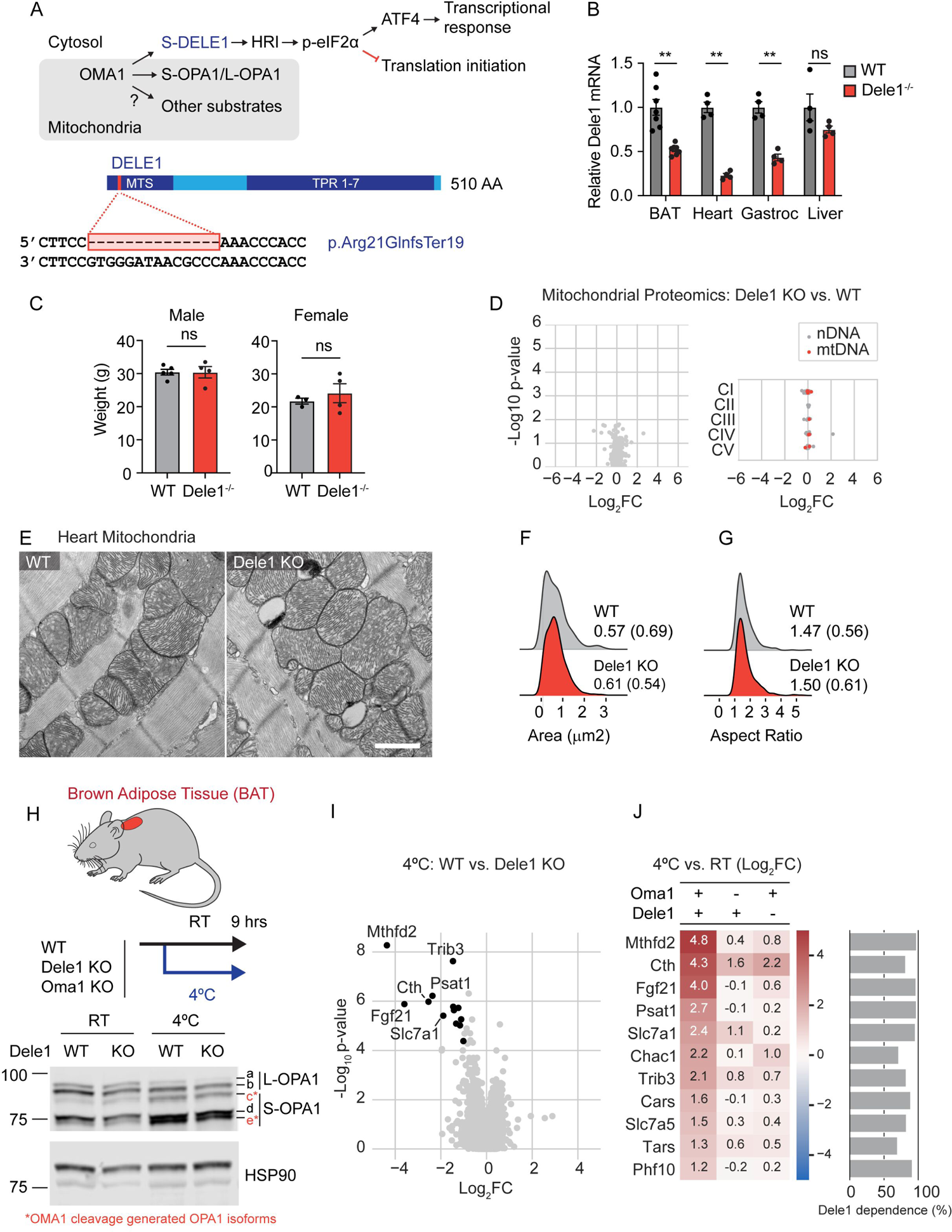
OMA1-DELE1 signaling mediates the integrated stress response under physiologic cold stress. (A) Diagram depicting the OMA1 stress signaling from mitochondria through cleavage of DELE1, activating the integrated stress response (ISR), and other substrates such as the inner mitochondrial membrane fusion protein OPA1 (top); and diagram of novel *Dele1* KO mouse, depicting the 510 amino acid (AA)-long DELE1 protein with mitochondrial targeting sequence (MTS) and tetratricopeptide repeat (TPR)-domains. The red bar indicates the site of 14 bp deletion in *Dele1*, resulting in a frameshift mutation (p.Arg21GInfsTer19) and early termination (bottom). (B) *Dele1* mRNA levels measured from microarray experiments of *Dele1* KO mice vs. WT littermates in four tissues indicated (BAT, brown adipose tissue; Gastroc, gastrocnemius muscle). Data from these microarray experiments also appears in (Fig. 3E, 5A-C, 5I, 7A-C). (C) Weights of Dele1 KO mice vs. WT littermates at 4 – 6 months of age. (D) Volcano plot represents all mitochondrial proteins (in MitoCarta3) measured from crude mitochondrial fraction of *Dele1* KO vs. WT littermates by mass spectrometry (left) and scatterplot depicting relative abundance of OXPHOS complexes I – V subunits. No mitochondrial proteins reached significance. WT group also is control for proteomics experiments in (Fig. 3F, 3I, 5G, and 5I). N = 4 mice per group. (E) Representative transmission electron micrographs of mitochondria from hearts of P28 *Dele1* KO and WT littermate mice. Scale bar = 1 μm. (F - G) Quantification of mitochondrial area and aspect ratio from TEM images like those shown in (E). Median values with interquartile range are shown. (H) *Oma1* KO, *Dele1* KO, and WT littermates (of *Dele1* KO) were challenged with cold stress for 9 hrs and interscapular brown adipose tissue was analyzed by immunoblotting for OPA1 cleavage by OMA1. OMA1 cleavage generated OPA1 isoforms are in indicated in red as c* and e*. (I) Volcano plot of global gene expression changes in cold stressed *Dele1* KO vs. WT littermates measured by microarray; significant genes (FDR < 0.05 and |Log_2_FC| > 1) are in black. N=7 mice per group. (J) Heat map depicting Log_2_FC for significant genes in (I) (left), and bar graph depicting the percent DELE1-dependence for the same genes (right).

To isolate the effects of OMA1 on the mt-ISR, we generated a constitutive *Dele1* KO mouse by CRISPR-Cas9 genome editing (Fig. 1A, bottom). No antibodies are currently available to detect endogenous DELE1, however, we verified that the mutation destabilizes *Dele1* mRNA in several tissues, including brown adipose tissue (BAT), heart, gastrocnemius skeletal muscle, and liver, likely by nonsense mediated decay (Fig. 1B). Additionally, the predicted truncated protein product (resulting from the predicted frameshift mutation p.Arg21GInfsTer19), if expressed, would lack part of the targeting sequence needed for mitochondrial localization and the tetratricopeptide repeat (TPR) domains required for HRI binding and downstream signaling (Yang *et al*, 2023).

*Dele1* KO mice appeared grossly normal with similar body weight to wildtype (WT) mice up to at least 4 – 6 months of life (Fig. 1C), and the heart mitochondrial proteome was not significantly altered by *Dele1* KO (Fig. 1D). Consistently, mitochondria from WT and *Dele1* KO hearts were indistinguishable by thin section transmission electron microscopy (TEM) at 28 days (Fig. 1E), with similar median areas (0.57 vs. 0.61 µm^2^) and aspect ratios (1.47 vs. 1.50) (Fig. 1F and G). Together, these findings suggest that *Dele1* KO does not disrupt mitochondria in the absence of mitochondrial stress.

We next considered whether DELE1 may be responsible for physiologic stress responses downstream of OMA1. We focused on cold stress in the heat-producing BAT, as cold stress has been shown to activate OMA1 and, in separate reports, cold stress has been reported to activate the ISR (Quirós *et al*, 2012; Jena *et al*, 2023; Flicker *et al*, 2019; Levy *et al*, 2023). Given OMA1 can be activated by mitochondrial uncouplers (Ehses *et al*, 2009; Head *et al*, 2009), activation of OMA1 in this setting is likely due to the fatty acid driven uncoupled mitochondrial respiration that produces heat (Nicholls, 2023). A 9-hour cold stress reduced BAT lipid droplets in WT, *Oma1* KO, and *Dele1* KO mice, as expected, indicating fatty acid consumption by mitochondria to support uncoupled respiration (Supplemental Fig. 1A). Additionally, OMA1 was activated to cleave L-OPA1 in WT and *Dele1* KO mice, producing the OMA1 specific cleavage products *c* and *e* from *a* and *b*, respectively (*d* is produced by cleavage of *a* by a different protease, YME1) (Fig. 1H) (Ehses *et al*, 2009; Head *et al*, 2009; Song *et al*, 2007). As expected, L-OPA1 cleavage was blocked by OMA1 KO (Supplemental Fig. 1B), confirming prior observations that OMA1 cleaves L-OPA1 in the setting of cold stress.

To determine whether DELE1 mediates the ISR, we assessed global gene expression in BAT. We reasoned that DELE1-dependent differentially expressed genes (DEGs) should change with stress in WT but not in *Dele1* KO mice. Intersecting the significant upregulated and downregulated DEGs from two comparisons (WT: stress vs. no stress, and Stress: *Dele1* WT vs. *Dele1* KO) (Supplemental Fig. 1C), we identified 11 DELE1-dependent DEGs, nearly all of which were genes classically associated with the ISR response, including *Mthfd2*, *Cth*, *Fgf21*, *Chac1*, *Psat1*, *Slc7a1*, *Slc7a5*, *Cars*, and *Tars* (Fig. 1I and J). For each gene, we also calculated the proportion of gene expression change that was attributable to DELE1 (% DELE1 dependence). This ranged from 70 - 97% for individual genes, indicating that DELE1 drives most of the gene expression changes in this set (Fig. 1J, right). Upregulation of these genes by stress was also largely dependent on OMA1, consistent with OMA1 functioning upstream of DELE1 in this pathway (Fig. 1J). As HRI mediates the ISR downstream of DELE1, this also explains why *Gcn2* KO in BAT did not block this response in a recent study (Levy *et al*, 2023). As expected, *Dele1* KO did not change gene expression in the non-stressed condition relative to WT (kept at room temperature) (Supplemental Fig. 1D). Thus, we identified the first DELE1-dependent physiologic stress: activation of the ISR in BAT in response to cold stress.

### DELE1 activates the mt-ISR in response to diverse mitochondrial stressors in striated muscle to promote growth and survival

We next asked whether DELE1 is responsible for the mt-ISR in response to diverse mitochondrial stressors. To address this question, we crossed the *Dele1* KO mouse with four different mouse models of mitochondrial myopathy/cardiomyopathy: two knock-in (KI) models of dominant mitochondrial myopathies *Chchd10*^G58R/WT^ (hereafter, C10 G58R) and *Chchd10*^S59L/WT^ (hereafter, C10 S59L) (Shammas *et al*, 2022; Genin *et al*, 2019; Anderson *et al*, 2019; Liu *et al*, 2020); and two KO models, *Chchd2*/*Chchd10* double KO (hereafter, C2/C10 DKO) and *Tfam* skeletal and cardiac muscle conditional knockout (*Tfam*^fl/fl^; *Ckmm*-Cre; hereafter, *Tfam* mKO) (Hansson *et al*, 2004; Liu *et al*, 2020; Nguyen *et al*, 2022) (Fig. 2A - L).

**Figure 2.**
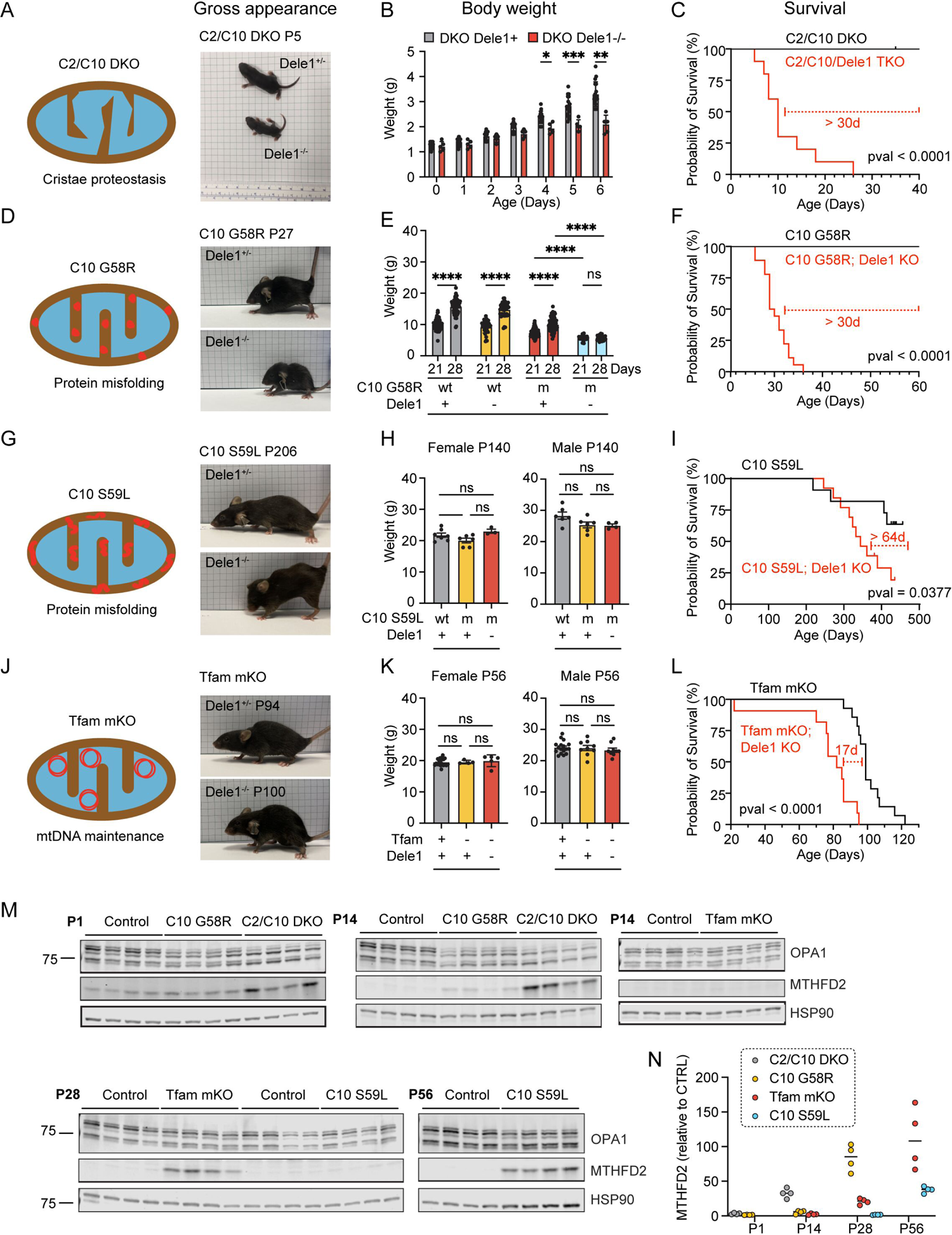
Dependence on DELE1 for growth and survival correlates with onset of mitochondrial stress in diverse models of myopathy and cardiomyopathy. (A) Cartoon depicting the cristae proteostasis stress induced by C2/C10 DKO and gross appearance of C2/C10 DKO mice with and without *Dele1* at P5. (B) Body weights of C2/C10 DKO mice with and without *Dele1* (*Dele1*+ indicates either *Dele1*^+/-^ or *Dele1*^+/+^) (C) Survival analysis of C2/C10 DKO mice. (D) Cartoon depicting protein misfolding stress of C10 G58R (aggregates in IMS) and gross appearance of C10 G58R mice with and without Dele1 at P27. (E) Body weights of C10 G58R mice with and without *Dele1* (*Dele1*+ indicates either *Dele1*^+/-^ or *Dele1*^+/+^). wt = wildtype; m = mutant. (F) Survival analysis of C10 G58R mice. (G) Cartoon depicting protein misfolding stress of C10 S59L (aggregates in IMS) and gross appearance of C10 S59L mice with and without Dele1 at P206. (H) Body weights of C10 S59L mice with and without *Dele1* (*Dele1*+ indicates either *Dele1*^+/-^ or *Dele1*^+/+^). wt = wildtype; m = mutant. (I) Survival analysis of C10 S59L mice. (J) Cartoon depicting disruption of mtDNA maintenance stress induced by *Tfam* mKO, and gross appearance of *Tfam* mKO mice with Dele1 at P94 or without Dele1 at P100. (K) Body weights of *Tfam* mKO mice with and without *Dele1* (*Dele1*+ indicates either *Dele1*^+/-^ or *Dele1*^+/+^). wt = wildtype; m = mutant. (L) Survival analysis of *Tfam* mKO mice. (M) Immunoblots of OPA1 cleavage and mt-ISR marker protein MTHFD2 expression, showing differential time course of activation of the mt-ISR postnatally, in the sequence C2/C10 DKO, C10 G58R, *Tfam* mKO, and C10 S59L. N = 4 mice per timepoint. Note: in wildtype mouse hearts MTHFD2 is expressed embryonically and on P1 but declines during postnatal development (Nilsson *et al*, 2014). (N) Quantification of MTHFD2 protein levels from immunoblots in (M). Levels for each group and timepoint are normalized to littermate controls except for C2/C10 DKO, which were normalized to controls for C10 G58R.

Importantly, the primary mitochondrial stress was different in each model. *Tfam* mKO is the prototypical model of OXPHOS deficiency due to a defect mtDNA maintenance (Hansson *et al*, 2004). This model is also predicted to induce severe mitonuclear imbalance, as expression of mtDNA encoded OXPHOS subunits is blocked but expression of nDNA encoded OXPHOS subunits is unaffected. C10 G58R and C10 S59L, by contrast, are models of intramitochondrial protein misfolding, and are not predicted to differentially effect levels of mtDNA and nDNA encoded OXPHOS subunits (Shammas *et al*, 2022; Genin *et al*, 2019; Anderson *et al*, 2019; Liu *et al*, 2020). The C10 G58R and S59L mutations cause CHCHD10 to misfold into two distinct toxic conformations within the mitochondria IMS with differential impacts on the mitochondria and the overall phenotype (Shammas *et al*, 2022). C10 G58R has an earlier and more severe impact on skeletal muscle than C10 S59L, leading to early-onset mitochondrial myopathy in both mice and humans (Shammas *et al*, 2022; Ajroud-Driss *et al*, 2015; Heiman-Patterson *et al*, 1997; Bannwarth *et al*, 2014; Anderson *et al*, 2019; Genin *et al*, 2019). C2/C10 DKO likely also disrupts mitochondrial cristae proteostasis but has a milder phenotype than protein misfolding from C10 G58R and C10 S59L (Liu *et al*, 2020; Huang *et al*, 2018). Indeed, the C2/C10 DKO are remarkable for having a near normal lifespan and healthspan, despite having early and pervasive activation of the mt-ISR (Liu *et al*, 2020; Nguyen *et al*, 2022).

Despite the diversity of the underlying stress, DELE1 was protective in each of the four models (Fig. 2C, F, I, and L). The survival benefit was especially pronounced for the C2/10 DKO and C10 G58R models (Fig. 2C and F), which also had the earliest activation of the mt-ISR, indicated by elevation of the maker protein MTHFD2 by postnatal day 14 (P14) (Fig. 2M and N). Strikingly, the C2/C10 DKO, which have a near normal lifespan and healthspan in the presence of DELE1 (Liu *et al*, 2020; Nguyen *et al*, 2022), lived a median of 10 days without DELE1 (Fig. 2C). Similarly, C10 G58R mice, which have a median life expectancy of more than 18 months with DELE1 (Shammas *et al*, 2022), survived a median of 1 month in the absence of DELE1 (Fig. 2F).

The early death in these models correlated with decreased growth in the first weeks of life. C2/C10/Dele1 triple KO mice had a similar weight as their siblings at birth, but their weight gain slowed from P4 (Fig. 2B). Similarly, weights of C10 G58R; *Dele1* KO mice were reduced relative to their littermates at 21 days and did not further increase from P21 to P28 (Fig. 2E). The failure to gain body mass also correlated with decreased motor function: C10 G58R; *Dele1* KO mice had decreased grip strength and an increased composite phenotype score compared to their C10 G58R; *Dele1*+ (either +/+ or +/-) siblings at 28 days (Supplemental Fig. 2A). Hand feeding only modestly increased survival of C10 G58R; *Dele1* KO mice (by 10 days), suggesting that food access was a minor contributor to their failure to thrive (Supplemental Fig. 2B). Hypertrophic growth of skeletal muscle accounts for about half of the 8-fold body mass increase in the first three weeks of life (White *et al*, 2010; Gokhin *et al*, 2008), and so we examined muscle fiber hypertrophy by measuring the cross-sectional area (CSA) of muscle fibers in the gastrocnemius muscle. Consistent with a decrease in skeletal muscle hypertrophy, body weights directly correlated with muscle fiber CSA among littermates in the C10 G58R; *Dele1* KO litters (r^2^ = 0.7272; Supplemental Fig. 2C). Thus, in the two myopathy/cardiomyopathy models with early DELE1 mt-ISR activation, early death correlated with decreased growth in the early weeks of life and worsening motor function.

In the other two models, *Tfam* mKO and C10 S59L, the mt-ISR was activated later: after P14 for *Tfam* mKO and after P28 for C10 S59L (Fig. 2M and N). The late activation of the mt-ISR in *Tfam* mKO mice likely reflects the postnatal expression of Cre from the *Ckmm* promoter, which has been estimated to recombine floxed alleles in heart and skeletal muscle between P7 and P21 (He *et al*, 2010). Late activation of the mt-ISR in C10 S59L mice is likewise consistent with our prior observations that C10 S59L reaches a higher protein abundance and takes longer to trigger an OMA1 stress response in both cultured cells and heart tissue compared to C10 G58R (Shammas *et al*, 2022). Body weights of *Tfam* mKO and C10 S59L were similar in the presence or absence of DELE1 (Fig. 2H and K). However, their heart to body weight ratio were significantly higher in the absence of DELE1 (Supplemental Fig. 2D and E), suggesting exacerbation of cardiomyopathy as the likely cause of early mortality in the absence of DELE1 (Fig. 2I and L).

Considered together, these findings suggest that the DELE1 mt-ISR is generally protective against diverse sources of mitochondrial stress in striated muscle and may be particularly critical when mitochondrial stress is present in striated muscle during early postnatal growth (Supplemental Fig. 2F).

### DELE1 functionally overlaps with OMA1 to protect against CHCHD10 myopathy

Next, we utilized the strong phenotype observed for C10 G58R; *Dele1* KO mice to genetically dissect the OMA1-DELE1 pathway. OMA1 has multiple substrates in addition to DELE1 in the mitochondria, including most notably the mitochondrial fusion protein OPA1 (Ehses *et al*, 2009; Head *et al*, 2009) (Fig. 1A). It is not known if the DELE1 mt-ISR confers all the survival benefit of the OMA1 stress response or whether cleavage of other substrates such as OPA1 also promotes survival. Additionally, recent cellular studies suggest that while OMA1 facilitates DELE1 signaling, in some settings DELE1 can signal the mt-ISR without cleavage by OMA1 (Sekine *et al*, 2023; Fessler *et al*, 2022). This would predict that *Dele1* KO may have a stronger effect on the mt-ISR than *Oma1* KO but this has not yet been tested *in vivo*. To address these questions, we directly compared the phenotypes of *Dele1* KO and *Oma1* KO mice under mitochondrial stress from C10 G58R protein misfolding, in litters triple mutant for C10 G58R, *Dele1*, and *Oma1* (Fig. 3A).

**Figure 3.**
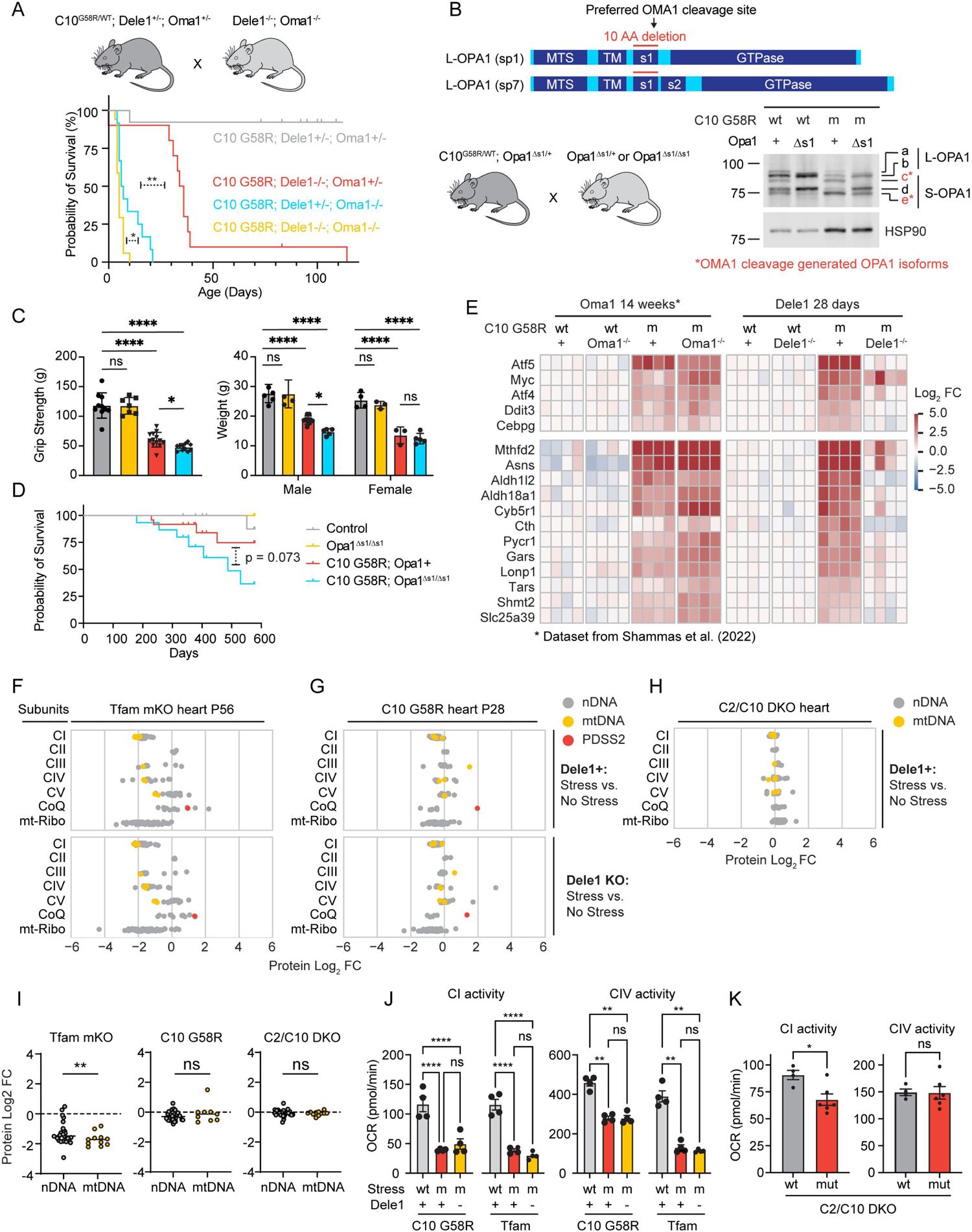
The DELE1 mt-ISR mediates only some of the protective OMA1 stress response and does not mitigate underlying OXPHOS complex dysfunction. (A) Illustration of breeding strategy to create the cohort. Survival analysis for C10 G58R; *Oma1* KO; *Dele1* KO triple mutant mice and their littermates. (B) Schematic demonstrating generation of *Opa1*^Δs1/Δs1^ mice through a 30 bp deletion removing the OMA1 preferred cleavage site, s1, from OPA1. Wildtype splice variants 1 and 7 (sp1 and sp7) contain the s1 site. (top); and OPA1 cleavage patterns of C10 G58R; *Opa1*^Δs1/Δs1^ mice and control mouse heart lysates (bottom, right). (C) Grip strength (left) and body weights (right) of C10 G58R; *Opa1*^Δs1/Δs1^ and their littermates at 13 weeks. Gray bar = control; yellow bar = *Opa1*^Δs1/Δs1^; red bar = C10 G58R; blue bar = C10 G58R; *Opa1*^Δs1/Δs1^. Each dot represents one mouse. (D) Survival analysis of C10 G58R; *Opa1*^Δs1/Δs1^ mice and their littermates. (E) Gene expression analysis of prespecified ISR genes from C10 G58R; *Dele1* KO mice and littermates at P28 compared to a previously published dataset from 14-week-old C10 G58R; *Oma1* KO mice (Shammas *et al*, 2022). Data from same C10 G58R; *Dele1* KO dataset also appears in (Fig. 1B, 5A-C, 5I, 7A). wt = wildtype; m = mutant. (F) Scatterplot depicting relative abundance of OXPHOS complexes I – V subunits, Coenzyme Q biosynthesis pathway, and mito-ribosome in *Tfam* mKO; *Dele1* KO mice and littermates detected from crude mitochondrial preparations of mouse hearts. See also Supplemental Fig. 3A for statistical analysis. N = 4 mice per group. Data from this proteomics dataset also appears in Fig. 5H and I. (G) Scatterplot depicting relative abundance of OXPHOS complexes I – V subunits, Coenzyme Q biosynthesis pathway, and mito-ribosome in C10 G58R; *Dele1* KO mice and littermates detected from crude mitochondrial preparations of mouse hearts. See also Supplemental Fig. 3A for statistical analysis. N = 4 mice per group. Data from this proteomics dataset also appears in Fig. 1D and 5G and I. (H) Scatterplot depicting relative abundance of OXPHOS complexes I – V subunits, Coenzyme Q biosynthesis pathway, and mito-ribosome in C2/C10 DKO and unrelated age-matched WT animals detected from crude mitochondrial preparations of mouse hearts. See also Supplemental Fig. 3A for statistical analysis. N = 4 mice per group. (I) Graphs compare protein abundance of mtDNA-vs. nDNA-encoded OXPHOS subunits from data in (F – H). (J and K) Seahorse oxygen consumption-based measurements of CI and CIV activities from frozen mitochondria isolated from hearts of the indicated genotypes. wt = wildtype; m or mut = mutant.

In these litters, *Oma1* KO had a substantially stronger effect on survival than *Dele1* KO (median survival 6.5 vs. 35 days), demonstrating that the *Oma1* stress response protects through multiple mechanisms, in addition to its activation of the DELE1 mt-ISR (Fig. 3A).

To identify additional mechanisms of OMA1 protection, we genetically blocked OMA1 from cleaving its other major substrate, OPA1, by generating a novel transgenic mouse line in which the preferred OMA1 cleavage site within OPA1, s1, is deleted (Ishihara *et al*, 2006) (Fig. 3B and Supplemental Fig. 2G). In primary fibroblasts from *Opa1*^Δs1/Δs1^ mice, the s1 deletion blocked basal cleavage and mitigated (but did not completely block) stress-induced cleavage by OMA1 (Supplemental Fig. 2G). The residual cleavage under stress is consistent with prior reports and suggests that OMA1 can cleave OPA1 at other sites in the absence of s1, albeit less efficiently (Ishihara *et al*, 2006). We next crossed *Opa1*^Δs1/Δs1^ and C10 G58R mice. Elimination of the s1 site reduced OMA1 cleavage of L-OPA1 both basally and in response to C10 G58R protein misfolding stress; cleavage of the *b* band to the *e* band was reduced by approximately 56% under basal conditions and 36% with C10 G58R stress, based on the e/b ratio (Fig. 3B). Partially blocking OPA1 cleavage in C10 G58R mice reduced body weight (significantly in male but not female mice) and grip strength at 13 weeks, suggesting that OPA1 cleavage by OMA1 is important for maintaining muscle function (Fig. 3C). However, the overall effect on C10 G58R mice was less pronounced than the effect of *Oma1* KO or *Dele1* KO, and the composite phenotype score and median survival were not significantly reduced (Fig. 3D and Supplemental Fig. 2H). The weak effect of the *Opa1* Δs1 allele is likely due to its incomplete block of OMA1 cleavage, although it is also possible OMA1 protects by cleaving other substrates in addition to OPA1 and DELE1. Nonetheless, these results suggest that cleavage of OPA1 by OMA1 is protective *in vivo*, independent of its activation of the mt-ISR.

We next considered whether DELE1 also retains some of its function in the absence of OMA1. We previously observed that approximately a quarter of C10 G58R; *Oma1* KO mice escape neonatal lethality and, surprisingly, have upregulation of mt-ISR-associated genes in the heart at 14 weeks (Shammas *et al*, 2022). This contrasted with knockdown of *Oma1* in adult C10 G58R animals, which reliably suppressed the mt-ISR associated genes (Shammas *et al*, 2022). Using gene expression analysis, we assessed whether the same genes could also be activated in the absence of DELE1. Notably, all C10 G58R; *Dele1* KO animals failed to activate this prespecified mt-ISR gene signature (Fig. 3E), including one outlier that survived to 16 weeks, close to the age of C10 G58R; *Oma1* KO mice with an activated ISR (Supplemental Fig. 2I). Consistent with the idea that DELE1 retains some limited function independent of OMA1, the C10 G58R; *Dele1* KO; *Oma1* KO triple mutants died earlier than the double mutant animals (Figure 3A, yellow vs. blue and red lines). Together these findings demonstrate that DELE1 is strictly required for activation of the mt-ISR in the setting of mitochondrial myopathy, in sharp contrast to OMA1. Thus, OMA1 and DELE1 have overlapping but separable protective effects on striated muscle *in vivo*.

### DELE1 mt-ISR has minor effects on OXPHOS complex subunits and mitochondrial structure

Next, to better understand how the DELE1 mt-ISR mediates protection against mitochondrial dysfunction, we considered whether the DELE1 mt-ISR corrects the underlying mitochondrial defect caused by stress in each myopathy/cardiomyopathy model.

The diversity of the models of myopathy/cardiomyopathy was reflected in the mitochondrial proteome and ETC complex activities resulting from C10 G58R protein misfolding stress vs. impaired mtDNA maintenance in the absence of TFAM (Fig. 3F – K and Supplemental Fig. 3A).

*Tfam* mKO, which causes decreased levels of mtDNA and mtDNA expression, resulted in the largest decrease in OXPHOS protein levels among the models (Fig. 3F, top, and Supplemental Fig. 3A). The average abundance of CI and CIV subunits was reduced to 31% and 43%, respectively, relative to controls (Fig. 3F, top). Consistently, CI, CIII, and CIV subunits were significantly enriched among MitoCarta3.0 Tier 3 pathways (Supplemental Fig. 3B). As expected, mtDNA-encoded subunits were significantly more downregulated than nDNA-encoded subunits in Tfam mKO hearts (Fig. 3F, gray vs. yellow data points; Fig. 3I), consistent with the primary defect in the expression of mtDNA-encoded subunits and mitonuclear protein imbalance. Mito-Ribosome and CoQ biosynthesis enzymes were also reduced, except for Pdss1 which is known to increase following disruption of CoQ biosynthesis pathway (Kühl *et al*, 2017).

C10 G58R, by contrast, led to a milder decrease in CI and CIV subunits (reduced to 72% and 93%, respectively, relative to controls) (Fig. 3G, top, and Supplemental Fig. 3A). The nDNA and mtDNA encoded subunits were similarly affected, suggesting that mtDNA expression was not limiting for OXPHOS protein expression and that C10 G58R does not induce a state of mitonuclear protein imbalance, in contrast to *Tfam* mKO (Fig. 3I). The OXPHOS complexes were least affected in adult C2/C10 DKO mice, which did not significantly decrease OXPHOS subunit expression (average CI and CIV subunit abundances were 91% and 107% of control levels, respectively) (Fig. 3H), and had comparatively mild effects on mitochondrial CI and CIV activities, with modestly decreased CI function (Fig. 3K).

Notably, *Dele1* KO did not significantly change OXPHOS subunit levels or CI or CIV activities in either C10 G58R or *Tfam* mKO models (Fig. 3F and G, bottom, Fig. 3J, and Supplemental Fig. 3A), suggesting that the underlying mitochondrial stress is distinct among models and is not resolved by the DELE1 mt-ISR.

We next examined the ultrastructure of heart mitochondria by thin section TEM. Mitochondria in TEM micrographs were analyzed using a combination of deep-learning segmentation and manual scoring of key morphological features in >600 mitochondria per genotype (8,923 mitochondria evaluated in total) (scheme depicted in Fig. 4A). As expected, the morphological defects were highly diverse among the four models, reflecting the diversity of the underlying mitochondrial stress. However, each model caused disruption of the cristae and the contiguous IMM, the site of both OXPHOS and OMA1 sensing of mitochondrial stress (Fig. 4B-F).

**Figure 4.**
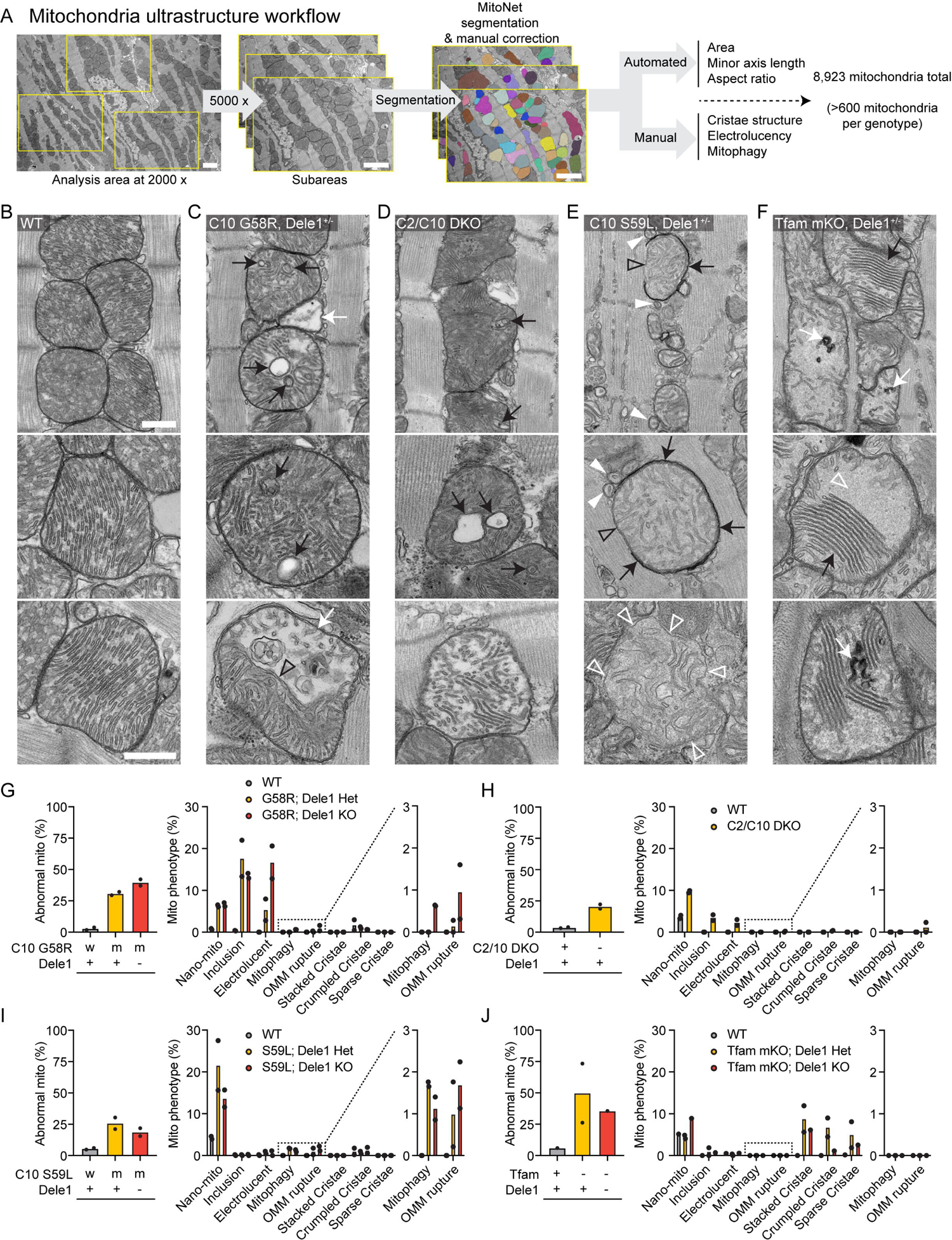
TEM shows that mitochondrial stressors cause diverse changes in mitochondrial structure, most of which are not suppressed by the DELE1 mt-ISR. (A) Workflow for analysis of mitochondria from hearts of diverse myopathy/cardiomyopathy models, combining deep learning aided segmentation of TEM images with manual scoring of characteristic ultrastructural features. Scale bars = 2 µm. (B) Representative TEM images of mitochondria from wildtype hearts. A field of view containing several mitochondria (top) and higher magnification views of individual mitochondria (below). Scale bars in B = 500 nm and apply to C-F. (C) Representative TEM images of mitochondria in C10 G58R hearts which exhibited cristal inclusions (black arrows) and electrolucent mitochondria (white arrow in top and bottom images). The bottom image shows a segmented type of electrolucent mitochondria with a cut-through cristae (open black arrowhead) forming the boundary between the electrolucent portion of the mitochondria, and that with a more typical appearance. (D) Representative TEM images of mitochondria in the C2/C10 DKO hearts which were characterized by cristal inclusions (black arrows) and the presence of electrolucent mitochondria (bottom). (E) Representative TEM images of mitochondria in C10 S59L hearts showed nano-sized mitochondria defined as measuring < 250 nm diameter in their minor axis with an aspect ratio < 1.5 (white arrowheads), mitochondria partially enclosed in electron-dense phagophore membranes (black arrows) with a portion of their periphery not enclosed (open black arrowheads), and mitochondria with a ruptured OMM (bottom image). Open white arrowheads indicate sites where the single layer of IMM is visible and OMM is absent. (F) Representative images of mitochondria in *Tfam* mKO hearts that displayed closely aligned stacked cristae (black arrows) and crumpled cristae (white arrows), as well as areas of where cristae were sparse and a homogeneous light grey matrix material was present (open white arrowheads). (G) Quantification of ultrastructural features for C10 G58R; *Dele1* KO at P28 and indicated littermates. (H) Quantification of ultrastructural features for C2/C10 DKO animals and unrelated age-matched controls at 3 – 4 months old. (I) Quantification of ultrastructural features for C10 S59L; *Dele1* KO at P140 and indicated littermates. (J) Quantification of ultrastructural features for *Tfam* mKO; *Dele1* KO at P56 and indicated littermates.

The C10 G58R model showed three prominent classes of abnormal mitochondria by thin section TEM: mitochondria with inclusions composed of cristae membranes (18% of mitochondria); nano-mitochondria, defined as measuring < 250 nm in their minor axis with an aspect ratio less than 1.5 (6% of mitochondria); and electrolucent mitochondria characterized by an enlarged matrix area absent of electron-dense substance and fewer cristae (5% of mitochondria) (Fig. 4C, G, and Supplemental Fig. 4), similar to the phenotype we observed previously (Shammas *et al*, 2022). The electrolucent mitochondria were sometimes segmented into two parts with an electrolucent portion separated by a cut-through cristae from an area of typical matrix and cristae density (Figure 4C, bottom and Supplemental Fig. 4H, bottom). These segmented mitochondria may originate from failed fusion between inner membranes, following successful fusion between outer membranes of electrolucent and normal mitochondria, as has been observed in the setting of *Opa1* KO (Hu *et al*, 2020). Consistent with this interpretation, the segmented mitochondria were approximately twice as large (in cross-section) as the unsegmented electrolucent mitochondria in our dataset (Supplemental Fig. 4B). Interestingly, mitochondrial ultrastructure in adult C2/C10 DKO hearts resembled that of C10 G58R, but with decreased frequency of cristal inclusions (4% of mitochondria) and electrolucent mitochondria (2% of mitochondria) (Fig. 4D, H, and Supplemental Fig. 5). This suggests that loss of C2/C10 function and C10 G58R misfolding may exert a similar stress on the IMM, which is more severe in the adult heart with C10 G58R misfolding.

Notably, similar proportions of nano-mitochondria and cristal inclusions were observed in C10 G58R mice in the presence or the absence of one copy of *Dele1* (Fig. 4G). Consistently, the average area of mitochondria was similar in the presence and absence of *Dele1* (Supplemental Fig. 4A). Electrolucent mitochondria were more frequent in the absence of *Dele1* (Fig. 4G), phenocopying a trend we observed previously for the effect of *Oma1* KO on electrolucent mitochondria (Shammas *et al*, 2022). These electrolucent mitochondria were also more likely to undergo outer membrane rupture and mitophagy, events that were observed almost exclusively in the absence of *Dele1* (Fig. 4G and Supplemental Fig. 4I and J). These results suggest that the DELE1 mt-ISR may be protective against the formation of electrolucent mitochondria in response to C10 G58R protein misfolding but did not affect other mitochondrial phenotypes, such as mitochondrial fragmentation and cristal inclusions, observed in C10 G58R mice.

In contrast to C10 G58R mice, the C10 S59L mice had few intracristal inclusions and electrolucent mitochondria, suggesting that C10 S59L exerts a different stress on mitochondria than C10 G58R, despite the proximity of the mutations (Fig. 4E, I, and Supplemental Fig. 6). Instead, the outer mitochondrial membrane (OMM) of C10 S59L mitochondria were more frequently ruptured and partially or fully enclosed by double-membraned autophagophores or autophagosomes, respectively, as has also been reported previously by others (Genin *et al*, 2019) (Fig. 4E, I, and Supplemental Fig. 6E and F). C10 S59L also had the highest frequency of nano-mitochondria (22%) in thin sections among the models (Fig. 4I). Tracking nano-mitochondria through several 60 nm-thick serial sections showed that some were indeed spherical, while others were short tubules (Supplemental Fig. 5G and H), but most were nano-tubes emerging from larger mitochondria and ending in a dead-end or connecting to other mitochondria. Similar numbers of OMM rupture and mitophagy events were observed in the presence or absence of *Dele1*, whereas fewer nano-mitochondria were observed in the absence of *Dele1* (Fig. 4I). The average area of mitochondria was similar in the presence and absence of *Dele1* (Supplemental Fig. 4A). Thus, the DELE1 mt-ISR did not prevent the mitochondrial morphological defects caused by C10 S59L protein misfolding stress.

Mitochondria from adult *Tfam* mKO hearts also had a distinct ultrastructure, compared to the other three models, suggesting loss of mtDNA exerts a different stress on mitochondria than C10 protein misfolding or C2/C10 DKO (Fig. 4F, J, and Supplemental Fig. 7). In *Tfam* mKO, mitochondrial cristae often appeared to adhere together forming patches of typically five or more “stacked” cristae, with little intralumenal space. Often, in the same mitochondrion, large regions of homogenous light gray matrix material were devoid of cristae, in a “sparse cristae” phenotype. Cristae that were crumpled into electron dense swirls were also observed, as well as cristae that appeared tubular, with increased diameter (Fig. 4F and Supplemental Fig. 7E). The extent of these phenotypes was highly variable among cardiomyocytes in the same sample, with mildly affected cardiomyocytes often bordering severely affected cardiomyocytes in a mosaic pattern. This mirrored the cellular heterogeneity observed by COX and SDH histochemistries and TEM in prior reports (Wang *et al*, 1999) (Supplemental Fig. 7B). Notably, these mitochondrial phenotypes were milder in a *Tfam* mKO; *Dele1* KO heart sample compared to two *Tfam* mKO; *Dele1*+ hearts, suggesting that the DELE1 mt-ISR likely does not protect against these mitochondrial defects (Fig. 4J).

Considered together, mitochondrial ultrastructure was distinct among the four models, although each resulted in disruption of the IMM. The DELE1 mt-ISR did not reverse the underlying mitochondrial phenotypes, excepting the electrolucent mitochondria in the C10 G58R model. Thus, the DELE1 mt-ISR does not mitigate most structural abnormalities caused by mitochondrial stress.

### The mt-ISR mediates a core transcriptional response to diverse mitochondrial stressors in striated muscle

To obtain a global view of the effect of the DELE1 mt-ISR on striated muscle, we performed transcriptomics in the hearts of three models that survived past weaning: C10 G58R, C10 S59L, and *Tfam* mKO. Despite the diversity of mitochondrial stress in these models, they shared 111 mito-stress DEGs in the heart, of which 51 (46%) were DELE1-dependent (hereafter, referred to as the DELE1 mt-ISR heart signature) (Fig. 5A – C and Supplemental Fig. 8A and B), with DELE1-dependency defined by the intersection of DEGS, as depicted in (Supplemental Fig. 1C). An additional 39 DELE1-dependent DEGs were shared in 2 of 3 models (Fig. 5B and Supplemental Fig. 8A). As an alternative approach to defined DELE1-dependent DEGs, we calculated the proportion of shared mito-stress DEGs that were at least 50% DELE1-dependent. This yielded a similar gene set of 57 genes (including all 51 DELE1 mt-ISR signature genes) (Supplemental Fig. 8C and D). Notably, the percent DELE1 dependence for genes in the DELE1 mt-ISR heart signature was similar regardless of the primary mitochondrial stress (Fig. 5D). Likewise, considering each model individually, a similar proportion of DEGs were DELE1-dependent (18 – 33%), indicating that the DELE1 mt-ISR accounts for a substantial portion of the overall transcriptional response to mitochondrial stress in each of the three models (Fig. 5E). Considered together, these results indicate that a common DELE1-dependent mechanism mediates signaling in response to diverse stressors such as mitochondrial protein unfolding and decreased mtDNA maintenance.

**Figure 5.**
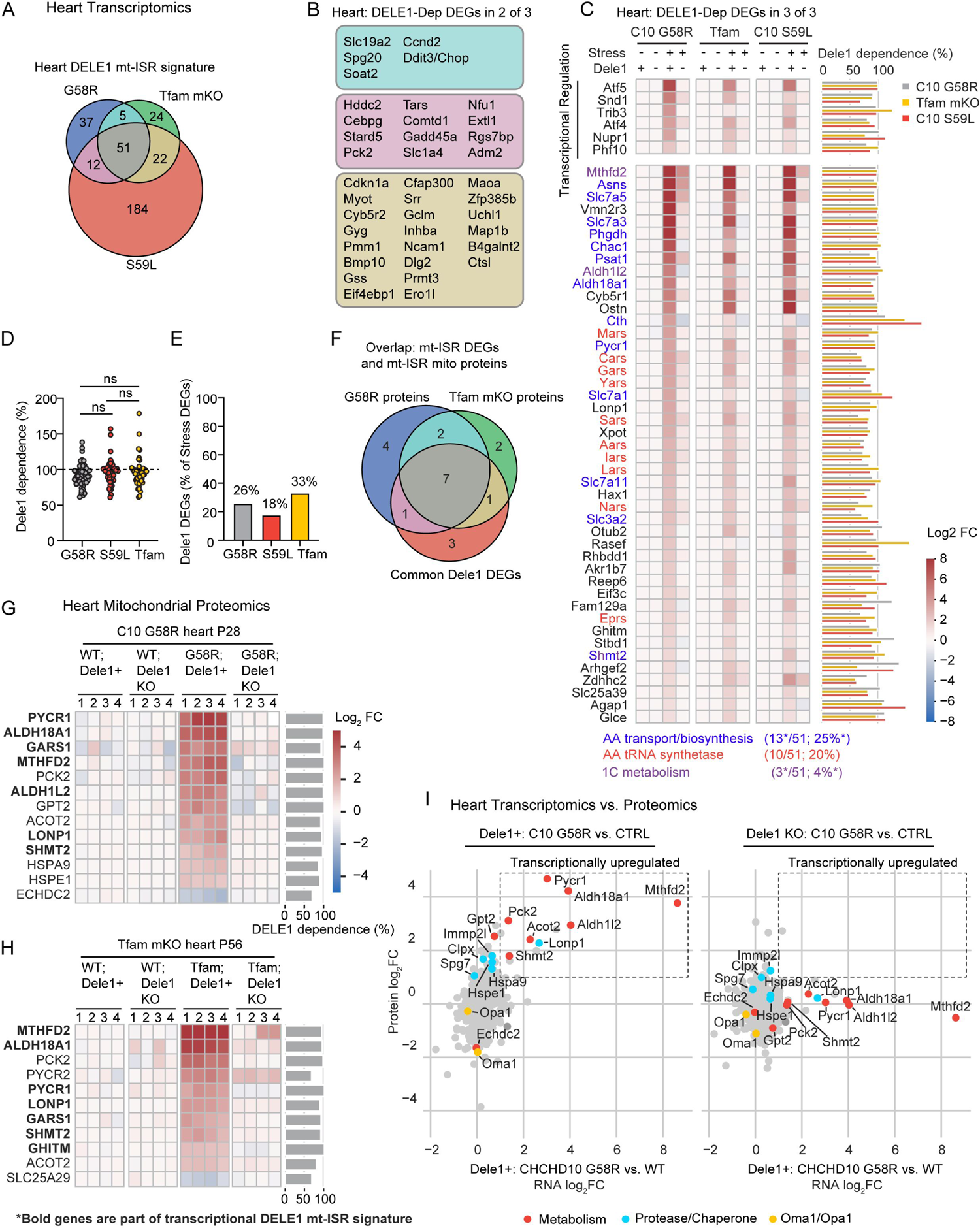
DELE1 drives a stereotyped transcriptional response to diverse mitochondrial stressors and reshapes the mitochondrial proteome in the heart. (A) Venn diagram depicting intersection of 51 DELE1-dependent DEGs common to all three different myopathy/cardiomyopathy models identified in microarray data used to define the Dele1 mtISR heart signature. DELE1-dependent DEGs are defined as in (Supplemental Fig. 1C). Data from the C10 G58R; *Dele1* KO dataset also appears in Fig. 1B, Fig. 3E, and Fig. 7A-B. (B) DELE1-dependent DEGs common to two of three myopathy/cardiomyopathy models (color coded as in A). (C) Heat map depicting fold change for DELE1-dependent DEGs identified in all three models (left). Those annotated as regulating transcription in (Lambert et al. Cell 2018) are separated from other genes. Bar graph on right depicts the percent DELE1 dependence for each gene. \**Shmt2* is in both the “AA transport/biosynthesis” and the “1C metabolism” categories. (D) Scatterplot compares percent DELE1 dependence for the 51 genes in the heart DELE1 mt-ISR signature among the three models. (E) Bar graph represents the percent of stress DEGs in each model that are significantly DELE1-dependent. (F) Venn diagram depicts intersection of the transcriptional DELE1 mt-ISR signature with significant DELE1-dependent changes in the mitochondrial proteome of C10 G58R and *Tfam* mKO animals. Some data from these proteomics datasets also appear in (Fig. 1D and Fig. 3F-I). (G) Heat map showing significant DELE1-dependent changes in mitochondrial protein abundance in C10 G58R hearts. N = 4 mice per condition. (H) Heat map showing significant DELE1-dependent changes in mitochondrial protein abundance in *Tfam* mKO hearts. N = 4 mice per condition. (I) Scatterplot compares RNA log_2_FC for C10 G58 vs. control (in the presence of DELE1) and mitochondrial protein log_2_FC for C10 G58 vs. control animals in the presence of DELE1 (left) or the absence of DELE1 (right).

Overall, the core DELE1 mt-ISR had several recognizable components, including: (1) transcriptional regulation (with *Atf5* having the highest fold-change in this group in all models); (2) amino acid transport and biosynthesis (particularly, of pathways for serine, glycine, asparagine, and proline biosynthesis) (13/51, 25% of genes); (3) protein translation, including amino acid tRNA synthetases, tRNA export (by *Xpot*), and protein synthesis re-initiation (by *Eif3c*) (12/51, 24% of genes) (Guan *et al*, 2017); and (4) mitochondrial 1C metabolism (which also intersects with serine and glycine metabolism and, through NADPH production, proline metabolism) (Ducker & Rabinowitz, 2017; Ducker *et al*, 2016) (Fig. 5C). Strikingly, nearly half of the genes in the core DELE1 mt-ISR signature were directed at promoting protein synthesis, as discussed further below.

These genes also overlapped extensively with those previously described as part of mitochondrial stress responses in mitochondrial myopathies, including *Atf4*, *Atf5*, *Mthfd2*, and *Lonp1*, all of which had > 81% DELE1-dependency in all three models (Fig. 5D) (Anderson *et al*, 2019; Tyynismaa *et al*, 2010; Dogan *et al*, 2014; Kühl *et al*, 2017; Nikkanen *et al*, 2016). Others previously associated with mitochondrial stress responses were either inconsistently elevated by mitochondrial stress or inconsistently DELE1-dependent. *Ddit3* (also known as, *Chop*), *Gdf15*, and *myc*, for instance, was strongly DELE1-dependent in some but not other models. The mitokine *Fgf21*, by contrast, showed a high DELE1 dependence (>83%) in all models but only increased > 2-fold in C10 G58R hearts. *Gpx4*, which was found to be regulated by the OMA1-DELE1 pathway in Cox10 cKO hearts (Ahola *et al*, 2022), was elevated by about 50% at the mRNA and protein levels across models tested but was not consistently DELE1-dependent (reaching nominal significance only in mitochondrial proteomics data from C10 G58R) (Supplemental Fig., 9A – D). Thus, DELE1 accounts for upregulation of many but not all classic markers of mitochondrial stress in mouse striated muscle, in response to diverse mitochondrial stressors.

Notably, a substantial proportion of the core DELE1 mt-ISR genes encode mitochondrial proteins (11/51, 21.6%). These were enriched for genes involved in metabolism (proline synthesis, serine/glycine biosynthesis, and 1C metabolism), in addition to three involved in quality control: *Lonp1*, encoding a matrix quality control AAA+ protease; *Ghitm* (also known as *Tmbib5*), recently identified to encode a H^+^/Ca^2+^ antiporter (Austin *et al*, 2022; Zhang *et al*, 2022; Patron *et al*, 2022); and *Slc25a39*, which was recently identified to encode a glutathione transporter (Wang *et al*, 2021; Shi *et al*, 2022). We compared this set to DELE1-dependent proteins in mitochondrial proteomes from C10 G58R and *Tfam* mKO mice (Fig. 5F). Notably, 6 of 7 detected mitochondrial proteins in this group were found to significantly increase with stress and were >91% DELE1-dependent at the protein level (Fig. 5G and H, proteins overlapping with Common DELE1 DEGs in **bold**). The exception was AKR1B7, which has multiple cellular locations in addition to mitochondria. Thus, DELE1-dependent changes in gene expression at the transcript level are largely reflected in the mitochondrial proteome in diverse models of mitochondrial stress.

In contrast to these DELE1 mt-ISR regulated genes, total mitochondrial protein levels of OPA1 and OMA1 were differentially regulated in the C10 G58R hearts and *Tfam* mKO hearts. Whereas OMA1 levels were decreased in C10 G58R hearts, consistent with its self-cleavage on activation (Fig. 5I), OMA1 levels were higher in *Tfam* mKO hearts than controls (Supplemental Fig. 9E and F). These results suggest that in the context of chronic stress, absolute levels of OMA1 and OPA1 are not reliable biomarkers for the OMA1-DELE1 mt-ISR activation, in contrast to ratio of OPA1 cleavage products and upregulation of DELE1-dependent genes.

The correspondence between transcriptional and protein level regulation for these core DELE1 mt-ISR genes prompted us to consider the inverse scenario: is the mitochondrial proteome transcriptionally regulated by stress independently of the DELE1 mt-ISR? Strikingly, we found that DELE1 accounted for all transcriptionally driven increases in the mitochondrial proteome of greater than 2-fold in C10 G58R hearts compared to control (Fig. 5I)). Similar results were obtained in *Tfam* mKO hearts, in which DELE1 accounted for all the transcriptionally driven increases in the mitochondrial proteome, with the exceptions of TIMM10, PRDX6, COMTD1, and MTHFD2 (Supplemental Fig. 9E and F). Notably, except for TIMM10 (in *Tfam* mKO) and IMMP2L (in C10 G58R), all increases in proteases or chaperones were either DELE1-dependent (as in the case of LONP1, HSPA9, and HSPE1) or were likely post-transcriptional (Supplemental Fig. 9G and H). This suggests that the DELE1 mt-ISR is the principal mito-nuclear signal mediating the mito-UPR in striated muscle.

### The DELE1 mt-ISR upregulates anabolic pathways that run through the mitochondrial matrix

Many of the amino acid biosynthesis pathways upregulated as part of the core DELE1 mt-ISR pass through the mitochondria, including those involved in asparagine synthesis (from the TCA intermediate oxaloacetate via aspartate), proline synthesis, and glycine synthesis (generated in the mitochondrial matrix from serine through the mitochondrial 1C metabolism pathway) (Fig. 6A, core genes in red). Flux through these interconnected pathways may be altered by OXPHOS dysfunction due, in part, to reductive stress in the mitochondrial matrix, as CI (and thus OXPHOS as a whole) is responsible for most of the NADH oxidation in the mitochondrial matrix. Consistent with reductive stress at the tissue level in adult C10 G58R hearts, NADP^+^ levels were significantly decreased and NAD^+^ levels trended toward decreased, in an untargeted metabolomic experiment (Fig. 6B).

**Figure 6.**
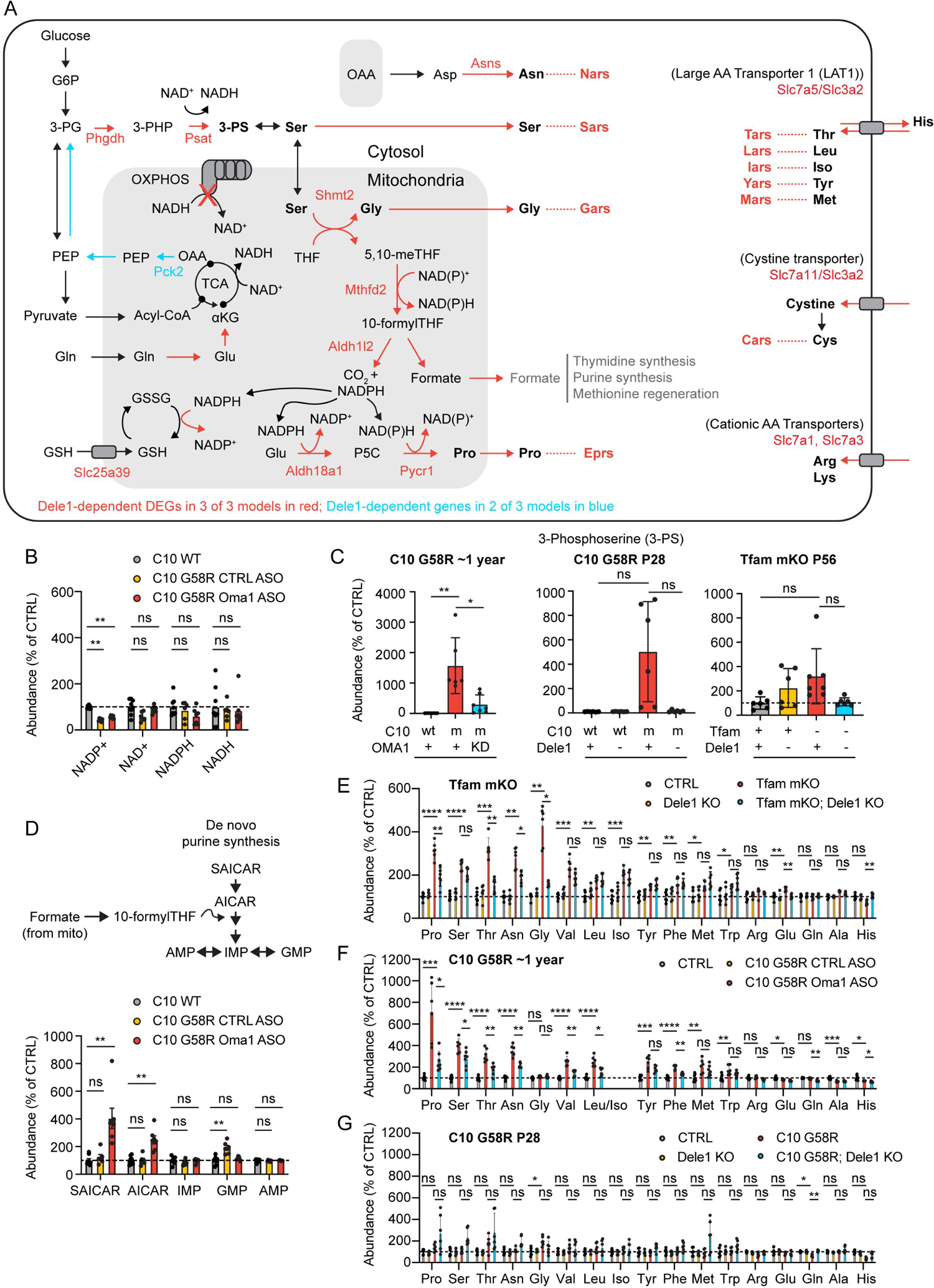
DELE1 mt-ISR maintains anabolic mitochondrial pathways, including for protein synthesis intermediates. (A) Diagram depicts major intersections among 22 (out of 51) proanabolic genes that are upregulated as part of the DELE1 mt-ISR heart signature (gene names in red). Additional genes (in blue) were identified in 2/3 models. Key pathways intersecting mitochondria include those for the biosynthesis of glycine, serine, proline, and asparagine. Upregulation of these pathways is coordinated with upregulation of genes for the corresponding aminoacyl tRNA synthases. (B) Levels of NADP^+^, NAD^+^, NADH, and NADPH detected by untargeted metabolomics of heart tissue from ∼1 year old (319 – 411 days) C10 G58R mice injected with CTRL or *Oma1* ASOs or wildtype littermates injected with PBS for 6 weeks prior to sacrifice. (C) Levels of 3-phosphoserine measured by untargeted proteomics in adult C10 G58R mice as in (B) or by targeted metabolomics in P28 C10 G58R mice and P56 *Tfam* mKO mice. W denotes wildtype; m denotes mutant. (D) Levels of intermediates of *de novo* purine synthesis that are sensitive to blocks in the mitochondrial 1C metabolism pathway measured from hearts of ∼1-year-old C10 G58R by untargeted metabolomics as in (B). (E-G) Levels of amino acids measured from hearts of P56 *Tfam* mKO, ∼1-year-old C10 G58R, or P28 C10 G58R mice as in (C).

We considered whether the OMA1-DELE1 mt-ISR may compensate for mitochondrial stress by upregulation of the key enzymes identified by transcriptomics and mitochondrial proteomics. To assess this in adult C10 G58R hearts, we used *Oma1* ASO knockdown strategy we employed previously (Shammas *et al*, 2022), as *Dele1* KO and *Oma1* KO cause early death in this model. Among the named metabolites in the untargeted metabolomics dataset, 3-phosphoserine (3-PS) had the greatest fold-change, increasing >100 fold in the hearts of C10 G58R mice compared to control mice (Fig. 6C and Supplemental Fig. 10A). This increase was blocked by *Oma1* KD, indicating it was dependent on the OMA1-DELE1 mt-ISR. This same pattern was also seen by targeted steady-state metabolomics in the hearts of P28 C10 G58R mice, and, to a lesser degree, in P56 *Tfam* mKO mice, although it did not reach significance in these models after correction for multiple comparisons (Fig. 6C and Supplemental Fig. 10B and C).

Notably, 3-PS is produced by two enzymes, PHDGH and PSAT1, the gene expression of which was strongly upregulated by the DELE1 mt-ISR in all models (Fig. 5C). These enzymes lie at a branch in glycolysis, in which glucose-derived carbons proceed to pyruvate, the TCA cycle, and ultimately OXPHOS or for the biosynthesis of serine and its derivates, such as glycine and 1C metabolism intermediates. The upregulation of PHDGH and PSAT1 and consequent increase in 3-PS are consistent with the hypothesis that a principal outcome of the OMA1-DELE1 mt-ISR is to shunt carbon from the glycolysis intermediate 3-PG toward biosynthesis.

Another potential carbon source for 3-PS synthesis is glutamine through the TCA cycle and gluconeogenesis. It is notable that two mitochondrial matrix enzymes, GPT2 (promoting glutamine entry into the TCA cycle via glutamate and oxaloacetate) and PCK2 (promoting gluconeogenesis from the TCA intermediate oxaloacetate) were increased by the DELE1 mt-ISR in C10 G58R hearts, and PCK2 was also increased in *Tfam* mKO hearts (Fig. 5B, G, H and Fig. 6A). Thus, the DELE1 mt-ISR may additionally push glutamine toward serine biosynthesis, as has been observed in rapidly dividing cancer cells with high biosynthetic needs (Vincent *et al*, 2015).

3-PS is the precursor for serine biosynthesis (Fig. 6A). Serine, in turn, is used for glycine biosynthesis and is the chief 1C donor for 1C metabolism (Ducker & Rabinowitz, 2017). Under basal conditions, most of the 1C metabolism flux from serine goes through the mitochondrial 1C metabolism pathway, with mitochondria exporting formate used for *de novo* purine and thymidine biosynthesis and the regeneration of methionine (after additional transformations) (Ducker *et al*, 2016). Mitochondrial 1C metabolism is also critical for generating the mitochondrial NAPDH pool needed for glutathione regeneration and proline synthesis, through the DELE1-responsive enzyme ALDH18A1. We, therefore, asked whether the OMA1-DELE1 mt-ISR helps maintain the mitochondrial 1C metabolism pathway in the setting of mitochondrial stress. To do so, we evaluated the levels of two metabolites AICAR and S-AICAR, precursors in purine synthesis, that are known to increase following blocks in mitochondrial 1C flux (particularly at MTHFD2) (Ducker *et al*, 2016). Consistent with the OMA1-DELE1 mt-ISR maintaining mitochondrial 1C flux in the setting mitochondrial stress, AICAR and S-AICAR levels remained baseline in C10 G58R mice but increased significantly when the OMA1-DELE1 mt-ISR was silenced, suggesting a partial block in the pathway in the absence of the mt-ISR (Fig. 6D). Thus, likely by upregulating mitochondrial 1C metabolism enzymes such as MTHFD2 and SHMT2 (and the upstream serine biosynthesis pathway), the OMA1-DELE1 mt-ISR may circumvent a block in the mitochondrial 1C metabolism pathway in the setting of mitochondrial stress.

In addition to serine and glycine biosynthesis, the DELE1 mt-ISR upregulates several amino acid biosynthesis and uptake pathways, including proline synthesis and asparagine biosynthesis, which connect directly with mitochondrial metabolism (Fig. 6A). We next evaluated whether these DELE1-dependent gene expression changes resulted in a steady-state change in amino acid levels in the hearts of adult C10 G58R (from untargeted metabolomics) and *Tfam* mKO mice (from targeted metabolomics). In both models, “aminoacyl-tRNA biosynthesis” was the most enriched KEGG metabolite set among mt-ISR dependent metabolites (Supplemental Fig. 10A and B). DELE1-dependent amino acids included proline, asparagine, and threonine in both models, and glycine, serine, leucine, phenylalanine, and glutamate in at least one of the models (Fig. 6E and F and Supplemental Fig. 10A and B). These DELE1-dependent amino acids all increased with stress.

We also examined amino acid levels in juvenile (P28) C10 G58R mice with one or no copies of *Dele1* (Fig 6G and Supplemental Fig. 10C). In contrast to the adult C10 G58R mice, there were no significant changes in the abundance of steady-state amino acids levels, except for glycine (which significantly increased with stress) and glutamine (which decreased with stress). The change in glutamine was significantly DELE1-dependent, whereas the change in glycine trended in the direction of DELE1 dependence but did not reach significance after correcting for multiple comparisons. The relatively small effect on the steady-state metabolome in the rapidly growing animal could be due to increased utilization of amino acids, particularly if amino acid abundance is one of the factors limiting growth in the C10 G58R animals. Interestingly, the C10 G58R; *Dele1* KO animals also showed high variability in steady-state tissue amino acid levels, which may reflect loss of homeostasis in animals that are at or nearing the end-stage of their life.

Considered together, these data suggest that the OMA1-DELE1 mt-ISR upregulates amino acid levels in the adult heart under mitochondrial stress, including four that are synthesized through pathways that are closely connected to mitochondrial metabolism: proline, asparagine, serine, and glycine. The steady-state increase of these four amino acids is likely driven by the DELE1 mt-ISR dependent upregulation of the enzymes responsible for their biosynthesis. That the upregulation of these amino acids is coordinated with the upregulation of their corresponding amino acid tRNA synthetase (i.e., proline with *Eprs*, asparagine with *Nars*, serine with *Sars*, glycine with *Gars*) additionally suggests that they may be upregulated to maintain adequate levels of tRNA-charged amino acids for protein synthesis (Fig. 5C and 6A).

### The DELE1 mt-ISR is similar in heart and skeletal muscle

We next asked whether the core transcriptional components for the DELE1 mt-ISR identified in the heart are conserved across mouse tissues. We compared the DELE1-dependent gene signatures among three mitochondrial enriched tissues from C10 G58R mice, the heart, the gastrocnemius, and the liver, as well as BAT from wildtype mice subjected to cold stress (Fig. 7A). Although OMA1 is activated strongly in the liver of C10 G58R mice (Shammas *et al*, 2022), the mt-ISR transcriptional response was surprisingly weak, with no DELE1-dependent DEGs at P28 and a small number of OMA1-dependent DEGs in tissue for C10 G58R mice evaluated at ∼1 year, following knockdown of *Oma1* with an ASO (Supplemental Fig. 11). Altogether, two OMA1 or DELE1-dependent genes, *Fgf21* and *Psat1*, were shared among all tissues, suggesting that these may represent the most robust markers for the OMA1-DELE1 mt-ISR across mouse tissues (Fig. 7A).

**Figure 7.**
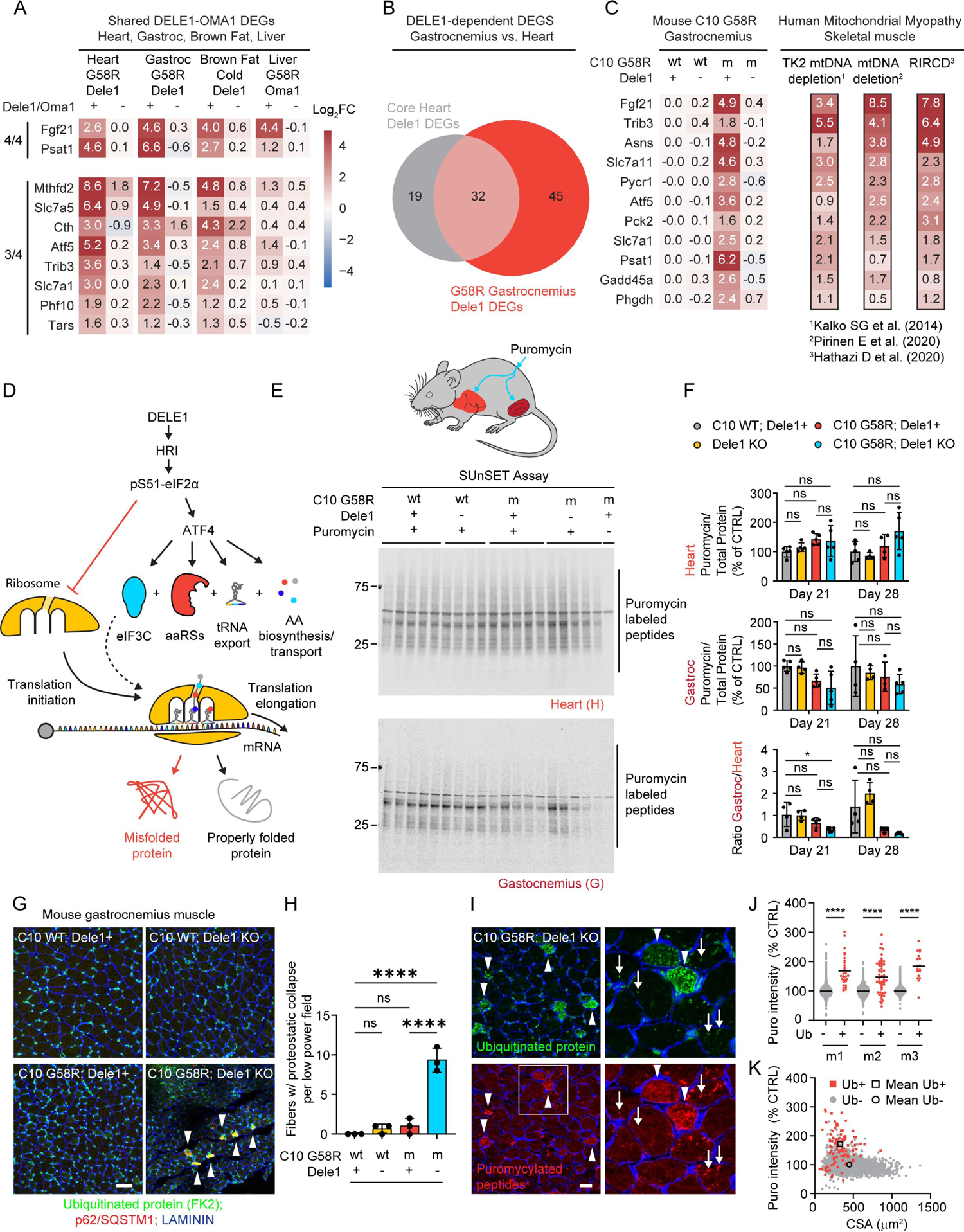
The DELE1 mt-ISR is overlapping in heart and skeletal muscle, where it does not decrease global protein synthesis but maintains proteostasis. (A) Heat map shows overlap among DELE1-dependent DEGS in four mitochondrial-rich tissues: heart, gastrocnemius (gastroc) skeletal muscle, and liver from C10 G58R mice, in addition to brown adipose tissue from mice subjected to cold stress as described in Fig. 1H. Brown adipose tissue microarray data from same dataset that also appears in Fig. 1B, 1I, and 1J. C10 G58R heart microarray data are from the same dataset that also appears in Fig. 1B, Fig. 3E, and Fig. 5A – B. C10 G58R liver microarray data are from the same dataset that also appears in Fig. 1B. (B) Venn diagram shows intersection of the heart DELE1 mt-ISR signature and C10 G58R DELE1-dependent DEGS from gastrocnemius skeletal muscle. (C) Heat map of 11 mouse genes that are DELE1-dependent in C10 G58R gastrocnemius model and have human orthologs that are upregulated by > two-fold (on average) in three previously published datasets from mitochondrial myopathy patients (Hathazi *et al*, 2020; Pirinen *et al*, 2020; Kalko *et al*, 2014). (D) Model depicts predicted effects of the DELE1 mt-ISR on protein translation including acute inhibition of translation initiation by pS51-eIF2α, resumption of protein initiation following the transcriptional upregulation of *Eif3c*, and facilitation of translation elongation through increased production of protein synthesis intermediates by the coordinated upregulation of (1) genes related amino biosynthesis and transport, (2) Xpot to mediate tRNA export from the nucleus to the cytosol, and (3) aminoacyl-tRNA synthases (aaRSs) to promote aminoacyl conjugation to tRNAs. Protein translation dynamics and fidelity can affect the proportion of newly translated proteins that fold properly vs. misfold. (E and F) Measurement of *in vivo* protein synthesis in C10 G58R; *Dele1* KO mice and littermates at P21 and P28, using the SUnSET assay. Mice were injected with the aminoacyl-tRNA ortholog, puromycin, sacrificed 30 min later, and puromycin incorporation into newly synthesized polypeptides in heart and gastrocnemius tissue lysates was measured by immunoblotting. (G) Representative immunofluorescence images of gastrocnemius muscle from P28 C10 G58R; Dele1 KO mice and littermates, showing fibers with many or confluent ubiquitinated protein aggregates co-localized with the aggregate-forming adaptor protein p62, suggesting proteostatic collapse (arrow heads) in low power (20X) images (top panels). Scale bars = 50 μm. (H) Quantification of (G). N = 3 mice per genotype with 30 low power fields were counted per genotype. Low-power field size was 397.75 μm X 397.75 μm. (I) Representative immunofluorescence images of gastrocnemius muscle from P28 C10 G58R; *Dele1* KO mice injected with puromycin 30 min prior to sacrifice as in (E). Muscle cross-sections were immunostained for ubiquitinated proteins (using the FK2 antibody) (green), puromycin (red), and LAMININ (blue). Arrow heads indicate muscle fibers containing many or confluent aggregates of ubiquitinated protein that were also co-stained for elevated puromycylated polypeptides. Arrows indicate individual aggregates of ubiquitin protein that also contain puromycylated polypeptides. Scale bars = 20 μm. (J and K) Quantification of (I). Individual myofibers were automatically segmented using the LAMININ immunofluorescence to define the myofiber border. Average puromycin fluorescence intensity and cross-sectional area were measured for each myofiber. Separately myofibers were manually scored as containing many or confluent ubiquitinated protein aggregates (Ub+ or Ub-muscle fibers, respectively). The average puromycin intensity for Ub+ and Ub-myofiber is shown in graph separately for three mice (m1 – 3) in graph. N = 3 mice with 10 high power fields counted per mouse. The scatterplot in (K) shows the relationship between myofiber cross-sectional area (CSA) and puromycin intensity for all myofibers analyzed in (J).

Among the mouse tissues, the heart and gastrocnemius showed the greatest overlap: 32 out of 51 (63%) mt-ISR heart signature genes were found in the gastrocnemius (Fig. 7B and C, and Supplemental Fig. 12A and B), including nearly all pro-anabolic genes diagrammed in (Fig. 6A).

Exceptions were *Iars*, *Cars*, *Mars*, *Eprs*, and *Shmt2*, which were strongly DELE1-dependent and trended toward being upregulated by C10 G58R stress but did not reach significance (Supplemental Fig. 12C). Interestingly, C10 G58R; *Dele1* KO mice also showed significant upregulation of catabolic genes, including those encoding the transcription factor *Foxo1a* and the atrogenes *Fbxo32* (also known as, *MAFbx*) and *Trim63* (also known as, *MuRF1*), suggesting that the DELE1 mt-ISR may suppress a catabolic program in skeletal muscle in addition to upregulating a pro-anabolic program (Supplemental Fig. 12D).

We next compared DELE1-dependent genes in skeletal muscle from the mouse C10 G58R mitochondrial myopathy model with the transcriptional signature of muscle from mitochondrial myopathy patients from three previously published datasets (Hathazi *et al*, 2020; Pirinen *et al*, 2020; Kalko *et al*, 2014). The patients in these datasets were of different ages and had different causes of their mitochondrial myopathy, but 11 genes in the mouse DELE1-dependent signature were upregulated by at least 2-fold averaged across the three studies (Fig. 7C). These included genes involved in biosynthesis of serine (*PHGDH* and *PSAT1*), proline (*PYCR1*), and asparagine (*ASNS*), as well as gluconeogenesis (*PCK2*), which may facilitate additional carbon sources for serine biosynthesis, as discussed above. The signature additionally included the amino acid transporters, *SLC7A11* and *SLC7A1*, for cystine and positively charged amino acids, respectively, and the mitokine *FGF21*. This suggests that key components of the DELE1 mt-ISR promoting anabolism in the setting of mitochondrial stress are conserved to humans and upregulated by diverse causes of mitochondrial myopathies.

### The DELE1 mt-ISR maintains protein synthesis and proteostasis in striated muscle under mitochondrial stress

The findings so far suggest that the DELE1 mt-ISR upregulates pathways to maintain protein synthesis intermediates in the setting of mitochondrial stress. This would be predicted to promote net protein synthesis by facilitating translation elongation (Fig. 7D). However, acutely the ISR also limits the availability of the ternary complex required for translation initiation, which would be predicted to slow the net protein synthesis rate (Pakos-Zebrucka *et al*, 2016). While these two actions of the ISR may seem opposed, they have in common the promotion of well-regulated protein synthesis under metabolic stress. By simultaneously restricting the number of ribosomes that can initiate translation and increasing protein synthesis intermediates to maintain translation elongation, the mt-ISR may minimize ribosome stalling and/or mistranslation, thereby limiting translation associated protein misfolding (Stein & Frydman, 2019; Stein *et al*, 2022). This suggests that the overall effect of the mt-ISR in mitochondrial myopathy may not be to slow net protein translation *per se* but to maintain the quality of protein translation in the setting of mitochondrial stress.

To directly assess the net effect of the DELE1 mt-ISR on protein synthesis in striated muscle, we performed the SUnSET assay in the heart and gastrocnemius muscle of the C10 G58R mice on the *Dele1*+ or *Dele1* KO backgrounds. Mice in each group were injected with puromycin, a tRNA aminoacyl analog that is incorporated into the elongating polypeptide, 30 minutes prior to sacrifice, and puromycylated peptides were detected by immunoblotting (Schmidt *et al*, 2009) (Fig. 7E and F). In C10 G58R mice, protein synthesis trended towards an increase in the heart and a decrease in gastrocnemius but did not differ significantly from control mice at P21 and P28 (Fig. 7E and F). Importantly, although *Dele1* KO normalized pS51-eIF2α in C10 G58R mice, as expected (Supplemental Fig. 12E), it did not increase net protein synthesis in the heart or gastrocnemius (Fig. 7E and F). We also considered the ratio of protein synthesis in the gastrocnemius to the heart for individual mice, in order to control for sources of interindividual variability. Notably, the C10 G58R; *Dele1* KO had the lowest heart-to-gastrocnemius ratio among groups, which reached statistical significance in comparison to control animals at the day 21 timepoint (Fig. 7F, bottom graph). Together these data suggest that the DELE1 mt-ISR does not have a net inhibitory effect on protein synthesis in stressed striated muscle, despite elevated pS51-eIF2α, and trended toward maintaining protein synthesis in skeletal muscle.

Having found the DELE1 mt-ISR did not decrease the net protein synthesis rate, we next considered its effect on proteostasis in skeletal muscle, using confocal microscopy. Notably, C10 G58R; *Dele1* KO but not C10 G58R; *Dele1*+ gastrocnemius muscle contained aggregates of ubiquitinated proteins that co-localized with p62/SQSTM1, a ubiquitin binding protein that sequesters misfolded proteins and can function as an autophagy adaptor (Supplemental Fig. 13A and B). In some fibers from C10 G58R; *Dele1* KO, these aggregates were so numerous that they appeared confluent by diffraction-limited light microscopy, suggesting proteostatic collapse within the myofiber (Fig. 7G and H).

Newly translated proteins are particularly vulnerable to protein misfolding, and the rate of protein translation can affect protein misfolding. To determine if protein aggregation within myofibers is associated with local changes in protein synthesis, we additionally assessed the gastrocnemius of C10 G58R; *Dele1* KO mice that were injected with puromycin 30 min prior to sacrifice (Fig. 7I - K). Myofibers with ubiquitin aggregates were again observed in C10 G58R; *Dele1* KO mice but not the other genotypes (Supplemental Fig. 13C). Notably, the muscle fibers with numerous aggregates had higher protein synthesis rates than other fibers in the same tissue (with 171.2% higher puromycin intensity on average). These myofibers also had a significantly smaller cross-sectional area (CSA) (mean CSA: 330.0 vs. 446.9 μm^2^), suggesting that they have failed to grow or are undergoing atrophy. Interestingly, individual ubiquitin protein aggregates also co-localized with puromycylated polypeptides, suggesting that the ubiquitin aggregates may form near sites of active translation. The puromycin immunostaining was likely specific for puromycylated polypeptides, as no signal was present in a negative control C10 G58R; *Dele1* KO mouse not injected with puromycin (Supplemental Fig. 13C, right most panel). These findings suggest that the DELE1 mt-ISR is important for maintaining translation-associated proteostasis in striated muscle under mitochondrial stress.

Considered together, these findings suggest that the DELE1 mt-ISR reshapes the metabolic network in stressed striated muscle to promote anabolism, including the continued production of intermediates for protein synthesis. We speculate that loss of the DELE1 mt-ISR may lead to loss of protein translation control and underlie the observed proteostatic collapse, decreased growth, and decreased survival in in models of early-onset myopathy, such as IMMD.

## Discussion

By comparing four myopathy/cardiomyopathy models resulting from diverse mitochondrial stresses, including protein unfolding and mtDNA depletion, we identified that the DELE1 mt-ISR is the predominant mitonuclear signaling response to mitochondrial stress in striated muscle. Although the primary source of mitochondrial stress was distinct in each model, they converged on disruption of the IMM, a central component of the OXPHOS system, which was sensed by OMA1 to activate the mt-ISR through DELE1 (Fessler *et al*, 2020; Guo *et al*, 2020). As OXPHOS disruption can affect the biosynthetic function of mitochondria, through decreased turnover of redox equivalents such as NADH (Luengo *et al*, 2021), the integrity of the IMM may be an important predictor of disruptions in mitochondrial metabolism. By sensing disruption of the IMM, the OMA1-DELE1 pathway may anticipate these disruptions to mitochondrial metabolism.

Notably, the mt-ISR did not correct the underlying mitochondrial structural or respiratory chain defects but rather compensated for OXPHOS dysfunction to maintain biosynthesis in the setting of stress, particularly through the upregulation of pathways for the biosynthesis of aminoacyl-tRNAs. It is notable that the ISR in yeast is largely a homeostatic response, in which uncharged tRNAs are sensed by GCN2, which, in turn, limits translation initiation, and triggers a transcriptional response to increase the synthesis of aminoacyl-tRNAs (Postnikoff *et al*, 2017). In this context, the OMA1-DELE1-HRI may be thought of as a predictive homeostatic response; it anticipates a limitation of biosynthetic intermediates in the setting of diverse forms of mitochondrial stress and responds by upregulating metabolic pathways to maintain biosynthesis. This response may not slow net protein synthesis *per se* but may promote the overall translation fidelity by reducing the chance of ribosome stalling or mistranslation.

Consistently, we found that the DELE1 mt-ISR did not slow net protein synthesis in the stressed heart and gastrocnemius, and, in fact, showed a trend toward promoting net protein synthesis in the gastrocnemius. This is also consistent with recent data from the *Dars2* cKO model, in which the ISR only slowed translation transiently in the setting of an exaggerated ISR response to chronic mitochondrial stress in the heart but did not mediate a prolonged decrease in translation (Kaspar *et al*, 2021). It is tempting to speculate that loss of translational control following *Dele1* KO may also explain the proteostatic collapse we observed in the striated muscle of C10 G58R; *Dele1* KO mice, as ribosome stalling and mistranslation are known to promote protein misfolding (Stein & Frydman, 2019; Stein *et al*, 2022). Consistently, myofibers with the most severely disrupted proteostasis in C10 G58R; *Dele1* KO mice had the highest rates of translation.

The DELE1 mt-ISR was particularly critical for survival in two models, C2/10 DKO and C10 G58R, that experienced mitochondrial stress onset during the rapid period of growth in the first weeks of life. The survival deficit was most dramatic in C2/10 DKO mice. We speculate that the DELE1 mt-ISR is particularly important for maintaining the biosynthetic capacity of stressed striated muscle during periods of rapid hypertrophic muscle growth. It is notable in this context that demands for protein synthesis intermediates are especially high in the first weeks of life in the mouse, as hypertrophic muscle growth accounts for half of the eight-fold increase in body mass during the first three weeks of life (White *et al*, 2010; Gokhin *et al*, 2008). This may explain why many of the pathways upregulated by the DELE1 mt-ISR in postmitotic striated muscle (such as serine/glycine synthesis, mitochondrial 1C metabolism, and proline synthesis) are also frequently upregulated in rapidly dividing cancer cells (Westbrook *et al*, 2022; Ducker *et al*, 2016; Geeraerts *et al*, 2021; Nilsson *et al*, 2014). In both cases, there is a strong demand for protein synthesis intermediates to increase biomass. Although striated muscle may be able to meet this need without upregulation of these pathways under basal conditions, in the setting of mitochondrial dysfunction, metabolic pathways may need to be rebalanced to maintain an uninterrupted supply of biosynthetic intermediates.

The dramatic survival benefit observed in the C10 G58R and C2/10 DKO models contrasted with the more modest survival benefit seen in the *Tfam* mKO and C10 S59L models. We speculate that the DELE1 mt-ISR is less important for survival in these models because the biosynthetic demands in striated muscle of juvenile and adult mice are less than those of neonatal mice. These demands are still present in the adult, however, and the DELE1 mt-ISR may similarly protect the *Tfam* mKO and C10 S59L models by maintaining biosynthesis in the setting of mitochondrial stress. The biosynthetic needs in the adult heart may be related to protein synthesis or they may be related to intersecting metabolic pathways.

Glycine and cysteine, for instance, are important for glutathionine synthesis and redox homeostasis. In the *Cox10* cKO, the DELE1 mt-ISR has been suggested to be important for redox homeostasis and prevention of ferroptosis (Ahola *et al*, 2022). In this context, it is noteworthy that among the 51 highly conserved genes across the models were the transporter for cystine, *Slc7a11*, which tends to increase intracellular cysteine levels (which contributes to glutathione synthesis together with glycine and glutamate), and the mitochondrial glutathione transporter, *Slc25a39*, which may also help prevent damage to membranes from lipid peroxidation and suppress ferroptosis. Likewise, serine, a 1C donor, is important for the *de novo* synthesis of nucleotides and the production of mitochondrial NADPH through the mitochondrial 1C metabolism pathway. In some models of mtDNA maintenance disorders like the Twinkle Deletor mouse, nucleotide synthesis through the 1C metabolism pathway may be particularly critical (Nikkanen *et al*, 2016). Thus, although the transcriptional response is similar in each model, the metabolic flexibility afforded by increased levels of key enzymes such as the NAD^+^-dependent PSAT1 and MTFHD2, may serve different biosynthetic needs at different developmental stages, and this may be reflected in downstream differences in the metabolome and proteome among the models.

Notably, the DELE1 mt-ISR transcriptional signature identified here overlapped extensively with that in other models of disrupted mtDNA maintenance or expression. Indeed, a similar response, identified first in Twinkle mutant mice (Tyynismaa *et al*, 2010), has been documented with disruptions at each step of mtDNA maintenance and expression, including mtDNA replication (Twinkle KO), mtDNA maintenance (*Tfam* KO), mtDNA transcription (*Polrmt* KO), mtRNA stability and processing (*Lrpprc* KO), and mtRNA translation (*Mterf4* KO and *Dar2* KO) (Dogan *et al*, 2014; Kühl *et al*, 2017). Our results from the *Tfam* mKO model strongly suggests that the DELE1 mt-ISR mediates this response resulting from decreased mtDNA expression or maintenance, generally. Thus, it is likely that the DELE1 mt-ISR underlies a transcriptional signature that it is seen in striated muscle across diverse forms of mitochondrial myopathy. As mtDNA mutations and disorders of mitochondrial maintenance are the most common causes of primary mitochondrial disorders (Gorman *et al*, 2015), the DELE1 mt-ISR may be an important response in most forms of mitochondrial myopathy. Consistently, we found extensive overlap between the DELE1 mt-ISR signature in the skeletal muscle of the IMMD mouse model of myopathy and genes upregulated in three cohorts of patients with mitochondrial myopathy, which included patients with mutations in Twinkle, *TK2*, and mtDNA (Hathazi *et al*, 2020; Pirinen *et al*, 2020; Kalko *et al*, 2014). This signature included genes involved in serine, proline, and asparagine biosynthesis, as well as the FGF21, which encodes a secreted mitokine that is widely used as a biomarker in mitochondrial myopathies (Lehtonen *et al*, 2016). This also suggests that FGF21 levels may provide a reliable marker of disruption at the IMM, sensed by the OMA1-DELE1 pathway.

Together our findings demonstrate a stereotyped DELE1-dependent response to diverse forms of mitochondria stress in mitochondrial myopathy. These mitochondrial stressors likely converge on disruption of the IMM and IMM-dependent OXPHOS. The response circumvents disruptions to biosynthetic pathways coupled to OXPHOS by rebalancing the metabolic network through a coordinated transcriptional program. This response is particularly critical during periods of rapid growth when biosynthetic demands are high. These findings suggest that interventions to promote the biosynthetic functioning of stressed striated muscle may be protective and caution against inhibition of the ISR in mitochondrial myopathy.

## Materials and Methods

### Mouse models

Mice were maintained on a 12-hour light/12-hour dark cycle, with food and water provided ad libitum unless otherwise stated. The Opa1^Δs1/+^ and Dele1^+/-^ mice were generated using CRISPR/Cas9 endonuclease-mediated genome editing on a C57Bl6J background by the NHLBI Transgenic Core. The deletions were confirmed by Sanger sequencing (Eurofins). *Ckmm*-Cre (JAX stock #006475) and *Tfam*^fl/fl^ mice (JAX stock #026123) were obtained from Jackson laboratory (Brüning *et al*, 1998; Hamanaka *et al*, 2013). The generation of C10 G58R, C10 G59L, C2/C10 DKO, and OMA1 KO mice were described previously (Liu *et al*, 2020; Shammas *et al*, 2022; Quirós *et al*, 2012). All animal studies were approved by the Animal Care Use Committee at the NINDS, NIH intramural research program. Both genders were used in all studies.

### Nutritional support

Mice in the nutritional support group were weaned on P28 instead of P21. In addition to the regular chow on the racks, they received KMR milk replacer using a small syringe twice daily from Monday to Friday and once daily on Saturday and Sunday. Other dietary supplements, including plain soft chow, Dietgel 76A (ClearH_2_O), and bacon-flavored treats, were also provided on the cage floor ad libitum.

### Cell Culture

Primary neonatal fibroblasts were generated using methods described previously (Liu *et al*, 2020). Primary neonatal fibroblasts generated from *Opa1*^Δs1/Δs1^ mice and from the wildtype littermates were treated with 20 μM CCCP or DMSO vehicle only for 16 hours.

### Cold Experiments

Adult mice in the experimental group were transferred to individual plastic cages with pre-chilled water but without bedding or food in a 4 °C cold room while mice in the control group remain room temperature. After 9 hours of exposure, mice were anesthetized with isoflurane and euthanized by cervical dislocation. Interscapular BATs were harvested immediately.

### Behavioral tests

Mouse forelimb grip strength was assessed using a BIOSEB instrument with grid attachment (catalog EB1-BIO-GS3). Three grip strength measurements were taken per mouse with 15-second resting periods between the trials. Composite phenotype scores were assigned based on the protocol described previously (Guyenet *et al*, 2010). All tests were performed by the same tester, who was blinded to the genotypes of the mice.

### RNA microarray

RNA microarrays were performed and analyzed as described previously (Shammas *et al*, 2022). In brief, RNA was extracted from frozen mouse hearts, gastrocnemius muscles, livers, and BATs using the Direct-zol RNA Miniprep Kit (Zymo, catalog R2051), the RNeasy Fibrous Tissue Mini Kit (QIAGEN, catalog 74704), or the RNeasy Lipid Tissue Mini Kit (QIAGEN, catalog 74804). RNA expression was measured using the Clariom S Mouse microarray (Affymetrix) by the NHGRI Microarray Core. Transcriptome Analysis Console software (Affymetrix, version 4.0.1) was used to analyze the data with the default settings.

### Immunoblotting

Immunoblotting and densitometric measurements were performed as described previously (Liu *et al*, 2020). In brief, mouse frozen hearts, BATs, and fibroblasts were lysed in RIPA buffer (Cell Signaling Technology, catalog 9806) with Protease/Phosphatase Inhibitor Cocktail (Cell Signaling Technology, catalog 5872). A buffer containing 20 mM Tris pH 7.8, 137 mM NaCl, 2.7 mM KCl, 1 mM MgCl2, 1% Triton X-100, 10% glycerol, 1 mM ethylenediamine tetraacetic acid and 1 mM dithiothreitol was used instead to lyse mouse frozen skeletal muscle. OPA1 bands on the blot were quantified using FIJI software (Schindelin *et al*, 2012). All other protein bands and total protein stains (LI-COR Biosciences, catalog 926-11016) were quantified using Image Studio v5.2 (LI-COR Biosciences).

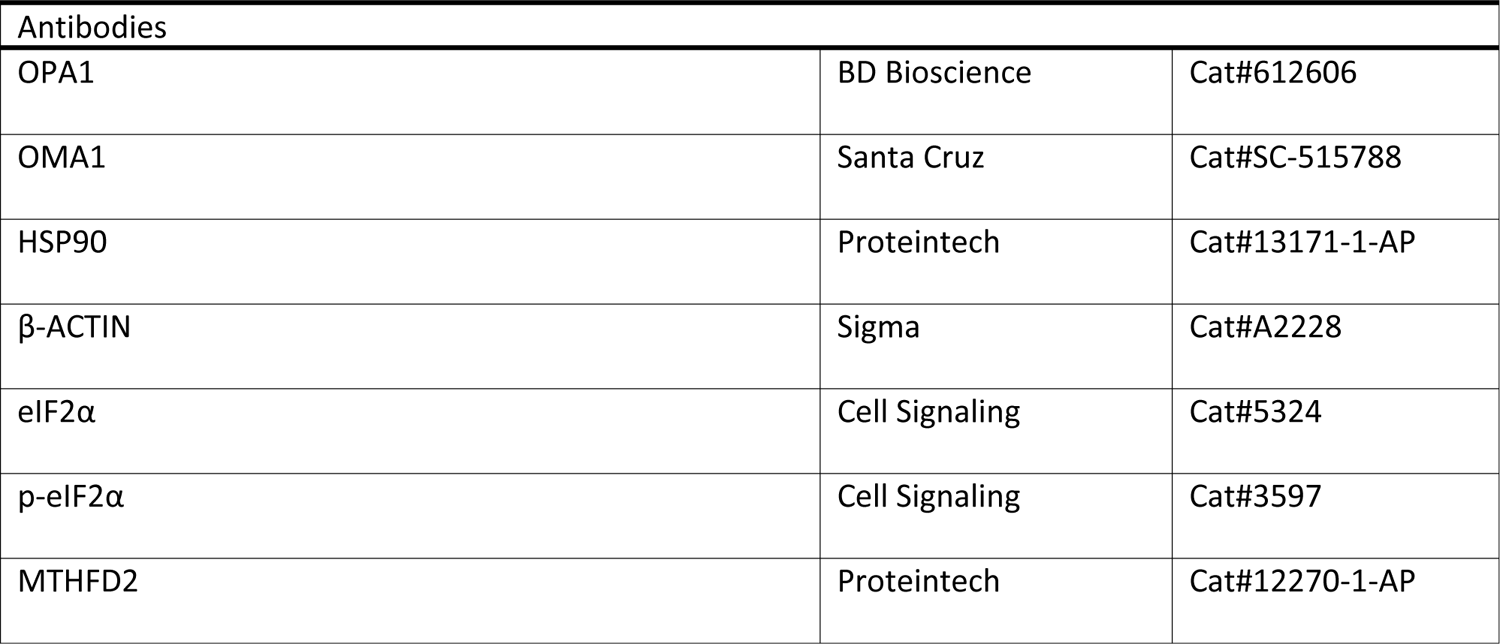

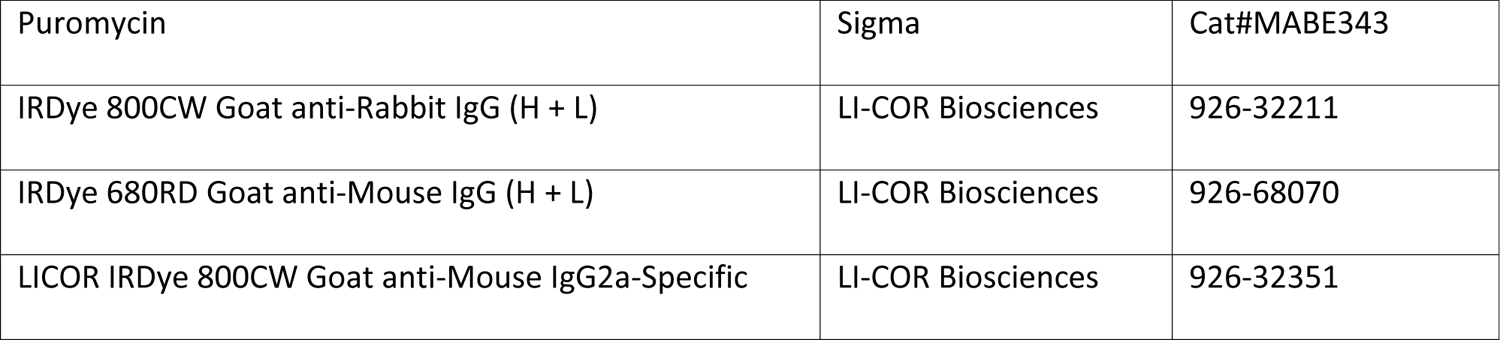

### Histological analysis

BATs were fixed in 4% paraformaldehyde in phosphate-buffered saline (PBS) overnight and washed in PBS three times. Fixed tissues were sent to Histoserv (Germantown, MD) for paraffin embedding, microtome sectioning, and haematoxylin and eosin (H&E) staining by standard procedures.

### Immunofluorescent Staining

Mice were anesthetized with isoflurane and transcardially perfused with PBS. Mouse tissues were dissected and frozen in chilled isopentane. 10 μm cross cryosections of the heart and of gastrocnemius mid-belly region were collected onto glass slides. Sections were washed with 0.1% Triton X-100 in PBS (PBST), blocked with 5% BSA in PBST for 1 hour at room temperature, and incubated in primary antibodies with 0.5% BSA in PBST overnight at 4 °C. Then, sections were washed in PBST and incubated in secondary antibody in PBST for 1-2 hours at room temperature. Slides were washed with PBST and PBS before coverslipped with ProLong Diamond Antifade Mountant (Invitrogen, catalog P36965) and sealed with nail polish. Images were obtained using Olympus FLUOVIEW FV3000 confocal laser scanning microscope.

| Antibodies |  |  |  |
| --- | --- | --- | --- |
| P62 | 1:200 | Abcam | Cat#AB56416 |
| Anti-ubiquitinated proteins antibody, clone FK2 | 1:200 | Sigma | Cat#04-263 |
| Laminin | 1:200 | Abcam | Cat#AB11575 |
| Puromycin | 1:500 | Sigma | Cat#MABE343 |
| 488 Goat anti-Mouse IgG1 | 1:500 | Invitrogen | Cat#A-21121 |
| 647 Goat anti-Mouse IgG2a | 1:500 | Invitrogen | Cat#A21241 |
| 555 Goat anti-Rabbit IgG (H+L) | 1:500 | Invitrogen | Cat#A32732 |

### Quantification of CSA of muscle fibers

Muscle fiber segmentation was performed using Cellpose 2.0 (Pachitariu & Stringer, 2022), deep learning program that utilizes human-in-the-loop training models to further automate the segmentation process, on laminin-labeled immunofluorescence images. After segmentation, CSA was quantified using LabelsToROIs of FIJI plugin (Waisman *et al*, 2021).

### Mitochondrial isolation

Mitochondria were isolated from mouse heart tissues as described previously (Frezza *et al*, 2007).

### Proteomics

Label-free quantitative proteomics of mitochondria isolated from mouse heart tissue was performed by the NINDS Proteomics Core Facility as described previously for most samples (Shammas *et al*, 2022). In brief, liquid chromatography-tandem mass spectrometry (LC-MS/MS) data acquisition was performed on an Orbitrap Lumos mass spectrometer (Thermo Fisher Scientific) coupled with a 3000 Ultimate high pressure liquid chromatography instrument (Thermo Fisher Scientific). Peptides were separated on an ES802 column (Thermo Fisher Scientific) with the mobile phase B (0.1% formic acid in acetonitrile) increasing from 3 to 22% over 70 minutes. The LC MS/MS data were acquired in data-dependent mode. For the survey scan, the mass range was 400-1500 m/z; the resolution was 120 k; the automatic gain control (AGC) value was 8e5. For C10 G58R and related genotype samples, a FAIMS interface was used. In this case the gradient MPB was increased from 3 to 20% over 63 minutes and the mass range of the MS1 scan was 375 - 1500 m/z. In all cases, the MS1 cycle time was set to 3 seconds. As many MS2 scans as possible were acquired within the cycle time. MS2 scans were acquired in ion-trap with an isolation window of 1.6 Da. Database search and mutant/WT ratio calculation were performed using Proteome Discovery 2.4 (Thermo Fisher Scientific) against Sprot Mouse database. Proteins were annotated as mitochondrial if they appeared in mouse MitoCarta3.0, and were additionally annotated with MitoPathways from MitoCarta3.0 (Rath *et al*, 2021).

### Seahorse assays

Seahorse oxygen consumption-based measurements of CI and CIV activities from frozen mitochondria isolated from hearts were performed according to the protocol described previously (Osto *et al*, 2020). 0.4 μg/well of frozen mitochondria were used with Seahorse XF Pro Analyzer (Agilent Technologies).

### ASO experiment

ASOs were synthesized at Ionis Pharmaceuticals as previously described (Seth *et al*, 2010; Shammas *et al*, 2022). Adult C10^wt/wt^; Oma1^+/–^ mice were given weekly subcutaneous injections of PBS. Adult C10^G58R/wt^; Oma1^+/–^ mice were given weekly subcutaneous injections of either a nontargeting (control) ASO or an OMA1-targeting ASOs. The mice were between 10.5 months and 13.5 months old. The ASOs were at a concentration of 5 mg/mL, and injections were dosed at 50 mg/kg. A total of 6 injections were administered per mouse over 6 weeks. Then, the mice were anesthetized with isoflurane and transcardially perfused with PBS, and their tissues were collected and flash-frozen in liquid nitrogen.

### Metabolomics

Mice were anesthetized with isoflurane and transcardially perfused with PBS. Mouse hearts were harvested and freeze-clamped immediately. Metabolomics data of the ASO experiment was obtained by the Metabolomics Core at the University of Michigan using their non-targeted metabolomics platform with manual peak integration on named compounds and automated peak detection for other mass spectrometer signals. Binner for data reduction and RefMet database (https://doi.org/10.1093/bioinformatics/btz798) were used (Kachman *et al*, 2020). Metabolomics data of the C10 G58R and Tfam mKO experiments were obtained by Metabolomics Core Resource Laboratory at NYU Langone Health.

### SUnSET *assays*

Measuring protein synthesis in mouse tissues was performed according to the protocol described previously (Ravi *et al*, 2020). In brief, puromycin 40 nmol/g of body weight was injected intraperitoneally to C10 G58R; Dele1 KO mice and their littermates. Mice were scarified after 30 minutes from the injection. Their tissues were collected and flash-frozen in liquid nitrogen. Immunoblotting was performed as described above.

### Transmission Electron Microscopy of Mouse Heart

Mice were deeply anaesthetized and transcardially perfused with PBS. Hearts were rapidly dissected, and sub-millimeter pieces of tissue were excised from the left ventricle and immersed in freshly prepared fixative containing 4% glutaraldehyde and 2 mM calcium chloride in 0.1M sodium cacodylate buffer, pH 7.4 (Electron Microscopy Sciences, Hatfield, PA, USA [EMS]). In the case of tissue obtained for the C2/C10 DKO mutant, mice were anaesthetized with isoflurane and then transcardially perfused with 2% paraformaldehyde, 2.5% glutaraldehyde, and 2 mM calcium chloride in 0.1M sodium cacodylate buffer. Perfusion-fixed hearts were excised and submerged in storage fixative containing 2% glutaraldehyde in 0.1M cacodylate buffer. Hearts were immediately further dissected by cutting a 1 mm-thick coronal section at the midpoint of the heart using a mouse heart slicer matrix (Zivic Instruments, Pittsburgh, PA, USA). The wall of the left ventricle in the coronal slice was cut into submillimeter pieces with a razor blade, submerged in storage fixative and stored at 4°C until processing for EM.

For EM processing, pieces of tissue were rinsed with 0.1M sodium cacodylate buffer (cacodylate buffer) three times and then postfixed on ice with reduced osmium containing 1% osmium tetroxide (EMS) and 1% ferrocyanide (Fisher Scientific, Pittsburgh, PA, USA) in cacodylate buffer for one hour. After three, 5-minute rinses in cacodylate buffer, tissue was rinsed in 0.1N acetate buffer (pH 5.0-5.2) at room temperature and then en bloc stained overnight with 1% uranyl acetate (EMS) in acetate buffer at 4°C. Tissue was then rinsed in acetate buffer and dehydrated through a graded series of 10-minute ethanol rinses prior to infiltration in EmBED812 epoxy resin using the manufacturer’s hard resin formulation (EMS). Finally, tissue was embedded in flat embedding molds and resin was polymerized in a 60°C oven for 48 hours.

Ultrathin sections were cut to a thickness of 60-70 nm using an ultramicrotome (EM UC7, Leica Microsystems, Wetzlar, Germany) equipped with a diamond knife (DiATOME, Hatfield, PA, USA). Sections were picked up on formvar coated 200-mesh or 1 mm-slot copper grids (EMS), and post-stained with 3% Reynold’s lead citrate (EMS) for 1 minute. Sections were viewed using a JEOL1400 Flash transmission electron microscope (JEOL USA, Inc., Peabody, MA, USA) operated at 120 KV. Images were acquired using a 29 Mpix CMOS detector (Advanced Microscopy Techniques, AMT, Danvers, MA, USA). Images from serial sections were aligned using the TrakEM2 plugin of Fiji image analysis software (Schindelin *et al*, 2012) which is a version of the open-source image analysis software, Image J2 (Rueden *et al*, 2017). For display, linear adjustments to light and dark levels in greyscale images were made Adobe Photoshop 2023 and figures were prepared using Adobe Illustrator 2023 (Adobe Inc., San Jose, CA, USA).

### Semi-automated analysis of mitochondria size, shape, and ultrastructural features in TEM images

Mitochondria were auto-segmented using MitoNet, a deep learning segmentation model, operated with Python software, using the napari plungin, called empanda (Conrad & Narayan, 2023). Structural features of mitochondria in TEM images were analyzed from two mice of each genotype (except for the *Tfam* mKO for which only one littermate of control and *Tfam* mKO, Dele1 KO were available). To carry out the analysis, images of at least two areas of tissue for each littermate were recorded at 2000 x direct magnification which encompassed a 550 µm^2^ field of view (examples for each genotype are shown in Supplemental Fig. 4 - 7). These areas were selected based on being located towards the interior of the tissue away from edges cut during dissection, and areas where the plane of the thin section was semi-longitudinal with respect to the muscle fibers, rather than transversely oriented. Within these areas, images of at least five subareas were acquired at 5000 x direct magnification, each of which encompassed an 87 μm^2^ field of view (examples are shown in Supplemental Fig. 4 – 7). Two or more of these 5000 x images were randomly selected for structural analysis using MitoNet segmentation. For each image, labels were automatically assigned to mitochondria detected in the TEM image by empanada-napari. The segmentation labels were manually proofread and corrected as needed and then the set of labels for each image was used to log the size features of each mitochondria including area, major and minor axis lengths, and aspect ratio into Excel (Microsoft Corporation, Redmond WA, USA). A minimum of 600 mitochondria per genotype were analyzed. The same labels created in empanada-napari were used to tabulate ultrastructural features of individual mitochondria in the control and mutant genotypes. With labels overlayed on the TEM image, each mitochondrion was examined and manually scored as normal or abnormal, and structural features noted in the Excel file that also contained the shape features logged for each mitochondrion. The percent of mitochondria with various structural features per genotype were graphed using GraphPad Prism 10.1.0 (316) version for Windows (GraphPad Software, Boston, MA, USA, www.graphpad.com).

### Statistics

Statistical analyses were performed using Prism (GraphPad), the statsmodels 0.14.1 Python module (for metabolomics and proteomics analysis), or the Transcriptome Analysis Console (TAC) (for analysis of microarray gene expression data). To define stress-dependent changes, two-tailed t-tests with pooled variance were performed comparing the stress; Dele1+ group vs. the control group. Multiple testing was corrected for using the Benjamini/Hochberg False Discovery Rate (FDR) method. Significant changes were defined as having a fold-change of >= 2 or <= 2 and an FDR <= 5%. DELE1-dependent changes were defined as significant stress-induced changes that were also significantly changed in the comparison of the stress; *Dele1* KO vs. stress; *Dele1*+ (applying the same fold-change and FDR cutoffs) with a fold change in the opposite direction to the Dele1+ group vs. the control group, consistent with reversion toward control. Percent DELE1 dependence was defined as the percent decrease or increase for *Dele1* KO vs. stress; *Dele1*+ divided by the percent increase or decrease for Stress; Dele1+ vs control. Set analysis for metabolomics data was performed using MetaboAnalyst 6.0 (https://www.metaboanalyst.ca/MetaboAnalyst/). In all figures,*, **, ***, and **** correspond to pvalues of <= 0.05, <= 0.01, <= 0.001, <= 0.0001, respectively.

### Data availability

Microarray data will be deposited in the NCBI’s Gene Expression Omnibus (GEO) Database. Analyzed data is available in Supplemental Tables 1 - 3.

## Acknowledgements

We thank Sandra Lara, Dr. Jung-Hwa Tao-Cheng, and the NINDS EM Facility for technical assistance with TEM. We thank Dr. Abdel Elkahloun and the NHGRI/DIR Microarray Core for technical assistance with RNA expression studies. We thank the NINDS Proteomics Core Facility for the label-free quantitative proteomics data acquisition. We thank Dr. Chengyu Liu and the NHLBI Transgenic Core for assistance in generating transgenic mice. We thank Drs. Pedro M. Quirós and Carlos Otin Lopez for providing the *OMA1* KO mice. We thank Dr. James J. Faust (Evident) for technical assistance with confocal microscopy. We thank Maureen Kachman and the Michigan Regional Comprehensive Metabolomics Resource Core for help with metabolic experiments of adult C10 G58R mice treated with *Oma1* ASOs.

We thank Ionis Pharmaceuticals for providing control and *Oma1* targeted ASOs that were used in in vivo experiments. We thank Dr. Richard Youle for critical reading of the manuscript and insightful comments. The linked image (https://en.wikipedia.org/wiki/Laboratory_mouse#/media/File:Vector_diagram_of_laboratory_mouse_(black_and_white).svg) was used in a modified form in some of the figures and is covered under a CC BY-SA 4.0 license. This work was supported by the Intramural Research Program of the NINDS, National Institutes of Health.

## Supplemental Figure Legends

**Supplemental Figure 1.**
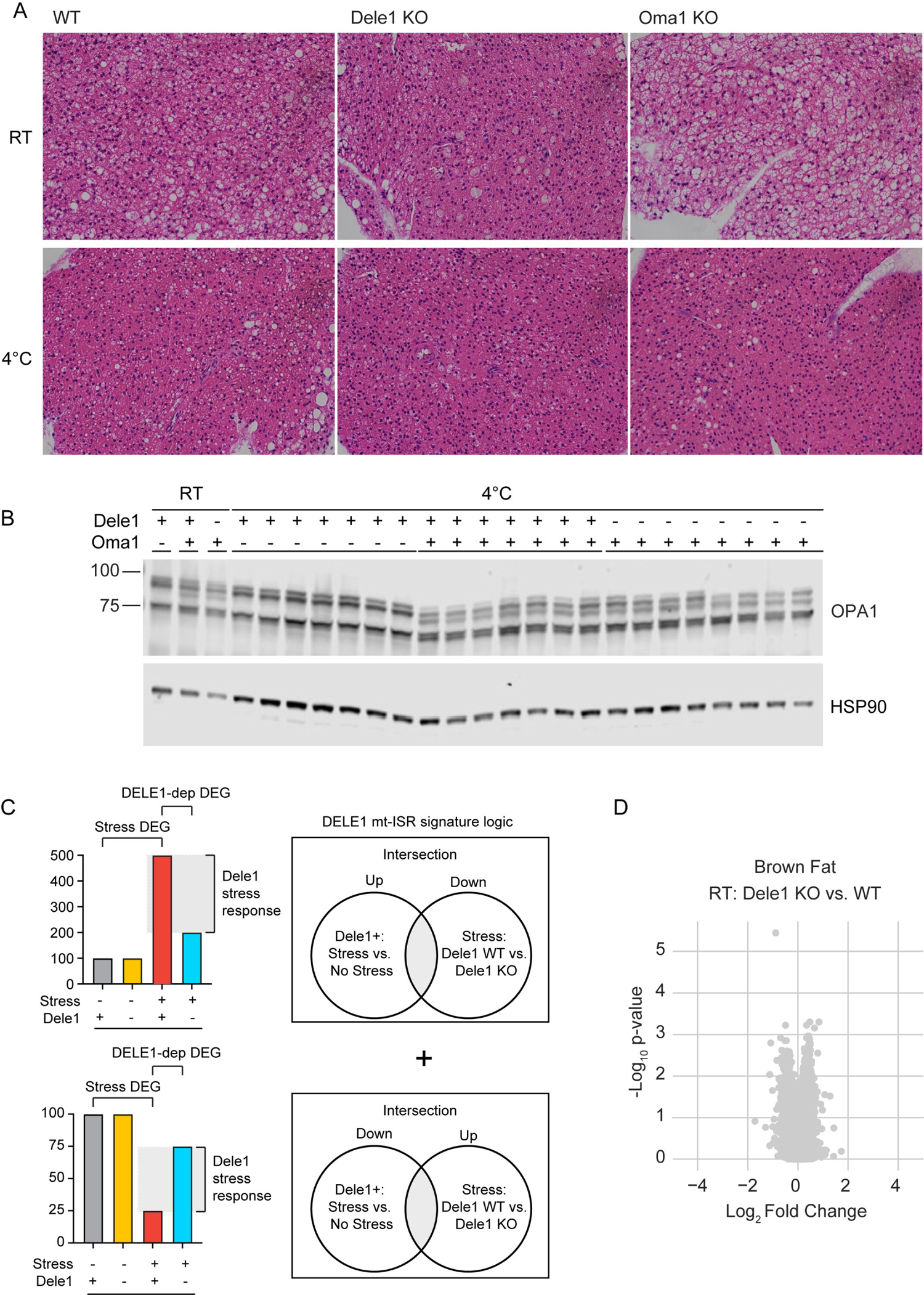
OMA1-DELE1 pathway mediates the integrated stress response in brown adipose tissue under cold stress. (A) H&E-stained section of brown adipose tissue from *Dele1* KO, *Oma1* KO, and WT littermates (of *Dele1* KO) subjected to cold stress in (Fig. 1H) shows reduction in brown adipose lipid droplets in all genotypes, which appear as unstained spheres, after cold stress. (B) Immunoblot showing OPA1 cleavage by OMA1 in WT and *Dele1* KO but not *Oma1* KO mice that were analyzed in (Fig. 1I). (C) Scheme depicts the logic used for defining DELE1-dependent DEGs from global gene expression data. (D) Volcano plot of microarray data from experiment in (Fig. 1I-J) comparing gene expression changes between *Dele1* KO and WT littermates left at room temperature (RT). No significant gene changes were observed. N=7 mice per group.

**Supplemental Figure 2.**
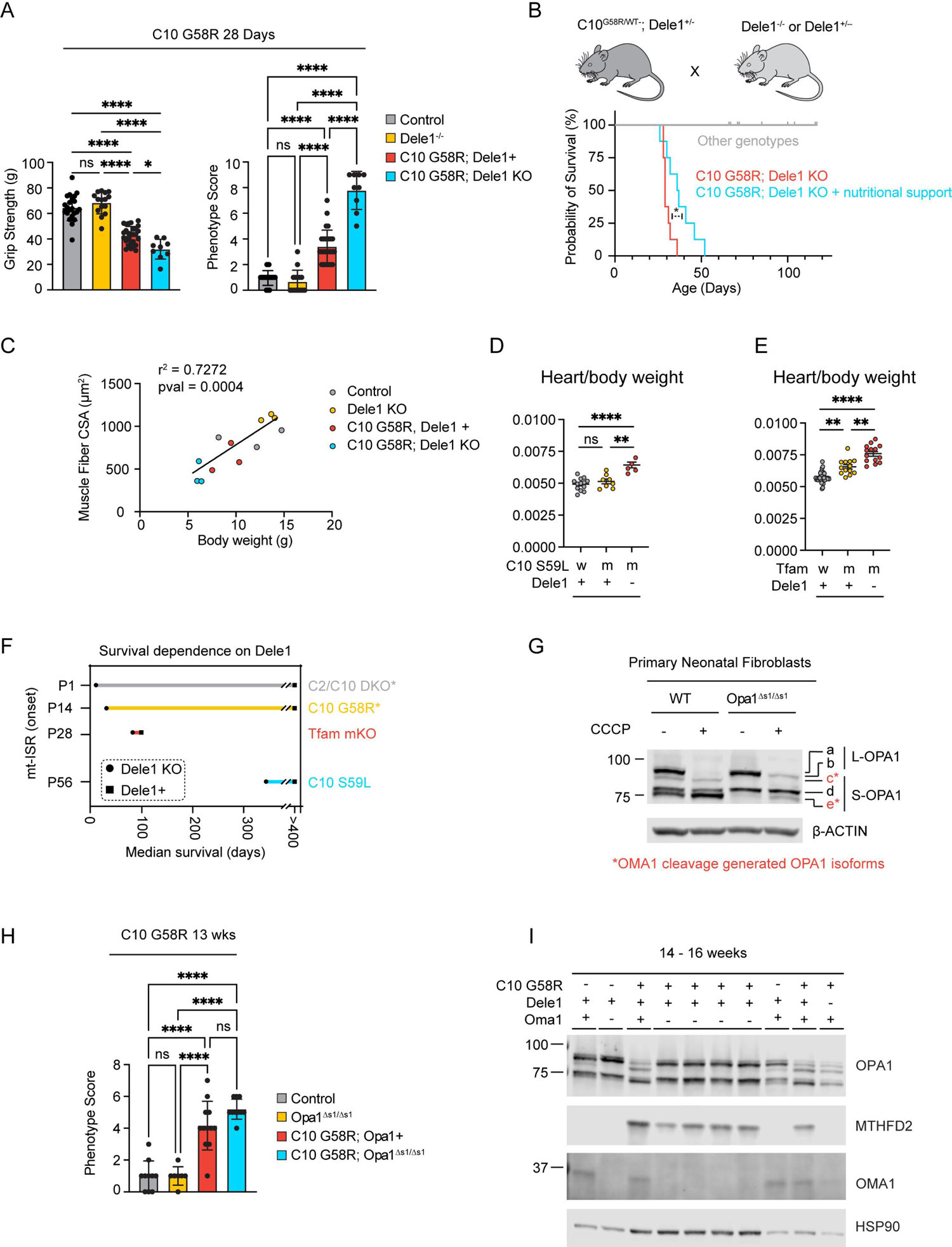
DELE1 mt-ISR promotes survival in diverse models of mitochondrial stress. (A) Grip strength (left) and a composite phenotype score (right) of C10 G58R; *Dele1* KO mice and littermates at P28. Composite phenotype score is comprised of ledge test, hindlimb clasping, gait test, and kyphosis. (B) Survival analysis of C10 G58R mice with and without nutritional support involving hand feeding twice per day. (C) Simple linear regression between gastrocnemius muscle fiber cross sectional area (CSA) and body weight for of C10 G58R; *Dele1* KO mice and littermates. (D) Heart to body weight ratio for C10 S59L mice; *Dele1* KO mice and littermates at P140. (E) Heart to body weight ratio for *Tfam* mKO mice; Dele1 KO mice and littermates at P56. (F) Correlation between age at stress onset and DELE1 survival benefit among the models. * indicates genotypes for which some lifespan estimates were determined from prior studies (Nguyen et al, 2022; Shammas et al, 2022). (G) Primary fibroblasts from *Opa1*^Δs1/Δs1^ mice treated with CCCP 20 μM or vehicle only for 16 hrs. The *c* and *e* bands generated by OMA1 cleavage (from *a* and *b*, respectively) are reduced at baseline and following uncoupling with CCCP; the *b* band is also relatively retained with CCCP, together demonstrating that L-OPA1^Δs1/Δs1^ (*a* and *b* bands) is resistant to OMA1 cleavage. (H) Composite phenotype score of C10 G58R; *Opa1*^Δs1/Δs1^ mice and littermates at 13 weeks. (I) Immunoblot compares OPA1 cleavage by OMA1 and elevation of the mt-ISR marker protein MTHFD2 in C10 G58R animals with or without OMA1 or DELE1 that survived to 14 – 16 weeks. Lysates from *Oma1* KO animals are from samples that were previously generated and appeared in (Shammas et al, 2022).

**Supplemental Figure 3.**
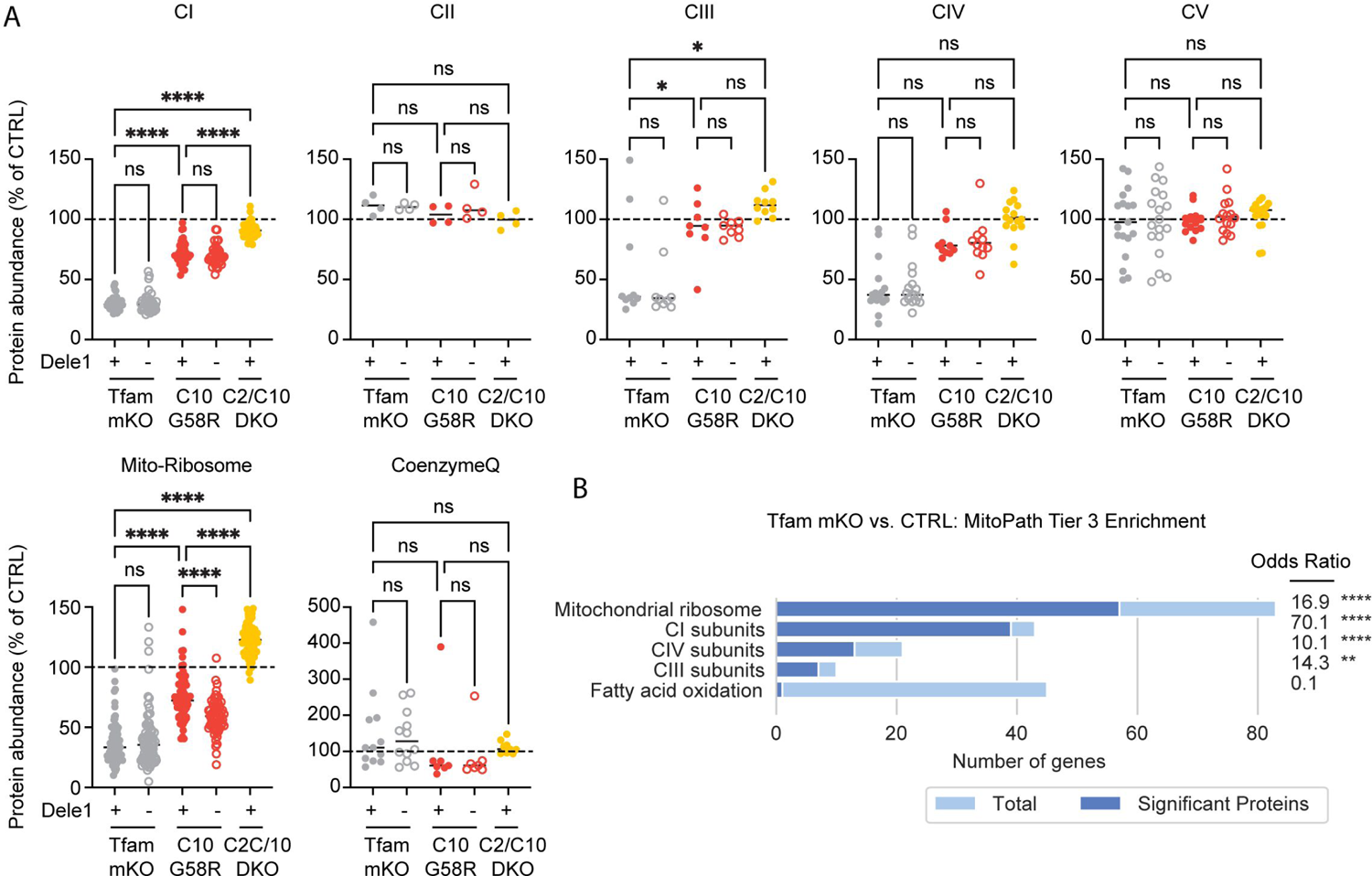
Comparison of OXPHOS subunit expression in heart mitochondria from diverse models of mitochondrial myopathy/cardiomyopathy. (A) Scatterplots depicting relative abundance of OXPHOS complexes I – V subunits, mito-ribosome, and Coenzyme Q from the indicated genotypes. Data are from the same datasets represented in (Fig. 3F – H), replotted to compare disease models. For statistics, a one-way ANOVA was performed followed by post-hoc testing, corrected for the multiple comparisons depicted within the graph with Dunnett’s test. All values are relative to littermate controls except for C2/C10 DKO which are matched to unrelated age-matched controls. Data from these proteomics datasets also appear in Fig. 5G - I. (B) Enrichment analysis mitochondrial proteins that significantly changed in *Tfam* mKO vs. control mitochondria isolated from hearts, using Tier 3 MitoPaths from MitoCarta 3.0.

**Supplemental Figure 4.**
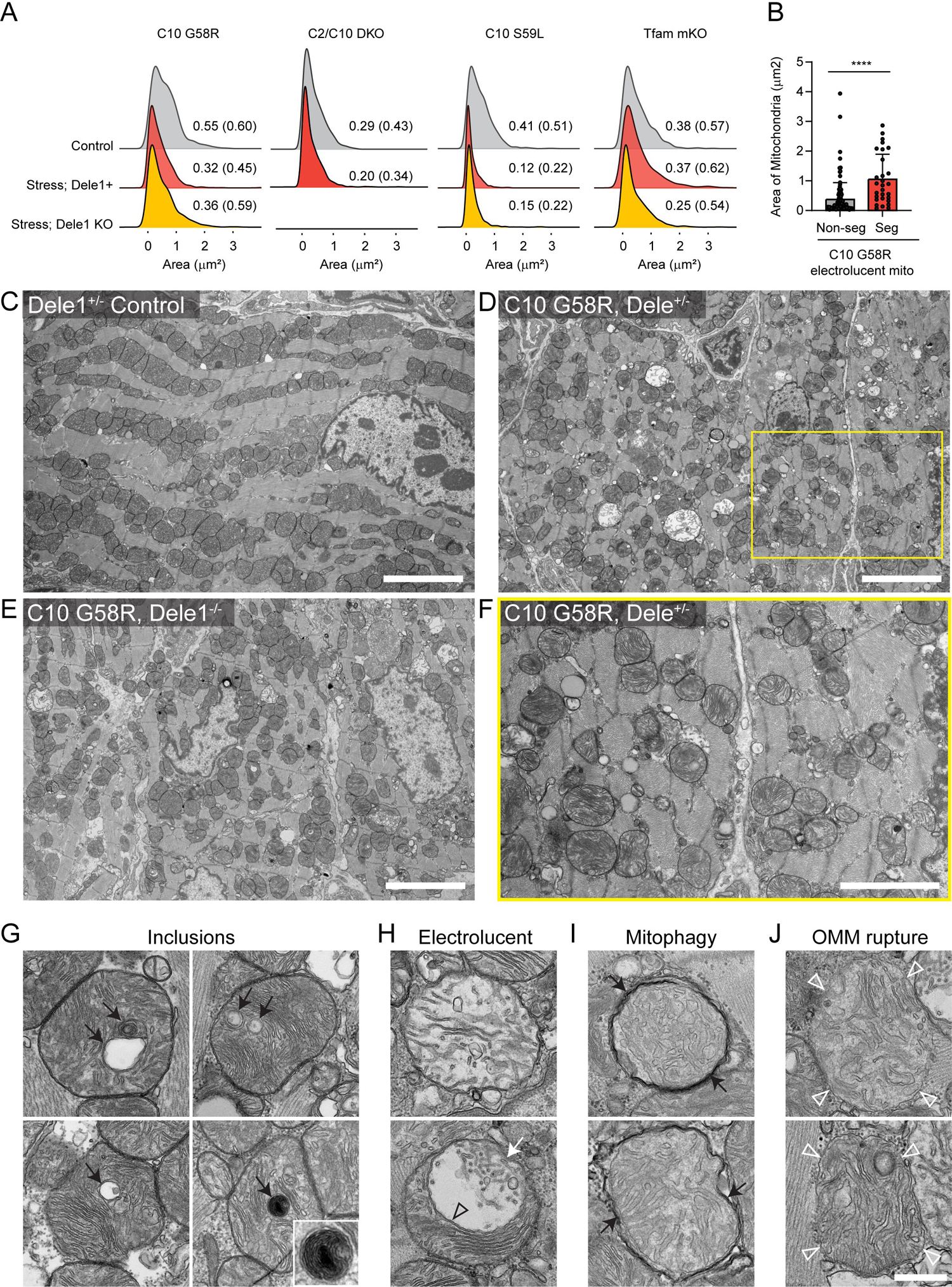
TEM of myocardium and ultrastructural features of mitochondria in C10 G58R on Dele1+/- and Dele1 KO backgrounds. (A) Kernel density plots showing distribution of mitochondrial areas for indicated genotypes, measured from TEM images of heart mitochondria. Median values and, in parentheses, interquartile ranges are reported adjacent to curves. N = 2 animals per genotype except for *Tfam* mKO; *Dele1* KO, where only 1 animal was available. > 600 mitochondria were measured per animal. (B) Bar graph comparing the areas of segmented and non-segmented types of electrolucent mitochondria that were obtained from analysis of C10 G58R animals and littermates in (A). Statistics were performed using Mann-Whitney test. (C - E) Representative TEM images acquired at 2000x direct magnification show areas of myocardium of indicated genotype used for analysis of mitochondria. Scale bar = 5 µm. (F) Image of the subarea boxed yellow in D, acquired at 5000x direct magnification and representative of the images used to quantify ultrastructural features of mitochondria detailed in Figure 4. Scale bar = 2.5 µm. (G) Examples of inclusions observed in C10 G58R mutant mitochondria (black arrows). (H) Examples of two types of electrolucent mitochondria characterized by an enlarged matrix area absent of electron-dense substance and fewer cristae. (Top) A uniformly electrolucent mitochondrion. (Bottom) a segmented mitochondrion that has an electrolucent part (white arrow) separated from a portion of normal-looking matrix and cristae by a cut-through cristae. Open black arrowhead indicates the junction between electrolucent and normal portions of the segmented mitochondria. (I) Mitochondria that are fully wrapped by electron-dense phagosome membranes (black arrows). (J) Mitochondria with ruptured OMMs. Open white arrowheads indicate sites where the intact IMM is visible, but OMM is absent. Scale bar in J = 500 nm and applies to G-J.

**Supplemental Figure 5.**
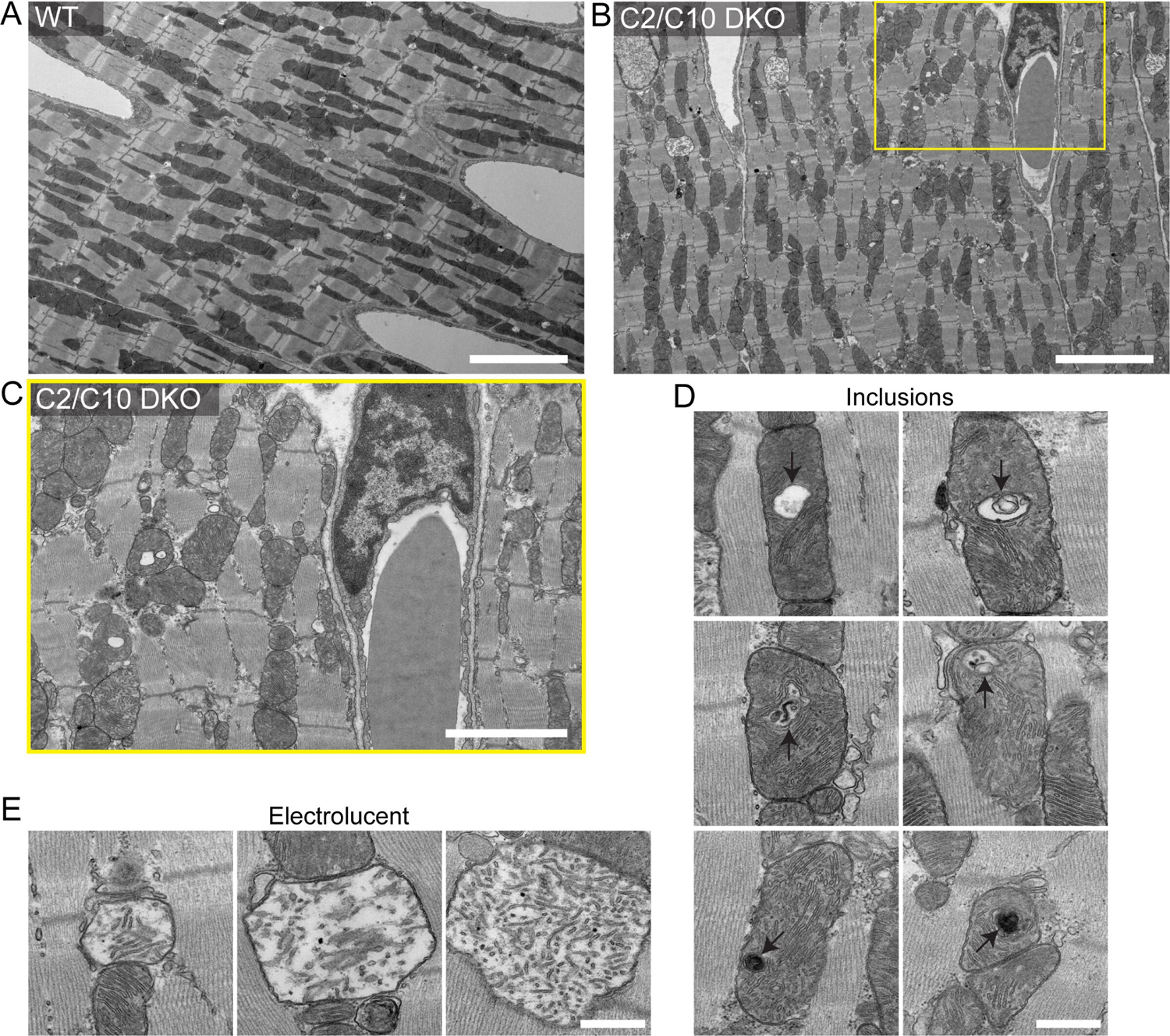
TEM of myocardium and ultrastructural features of mitochondria in C2/C10 DKO. (A-B) Representative TEM images acquired at 2000x direct magnification show areas of myocardium of indicated genotype used for analysis of mitochondria. Scale bar = 5 µm. (C) Image of the subarea boxed yellow in B, acquired at 5000x direct magnification and representative of the images used to quantify ultrastructural features of mitochondria detailed in Figure 4. Scale bar = 2.5 µm. (D) Examples of inclusions observed in C2/C10 DKO mitochondria (black arrows). Scale bar = 500 nm. (E) Examples of electrolucent mitochondria characterized by an enlarged matrix area absent of electron-dense substance and fewer cristae. Scale bar = 500 nm.

**Supplemental Figure 6.**
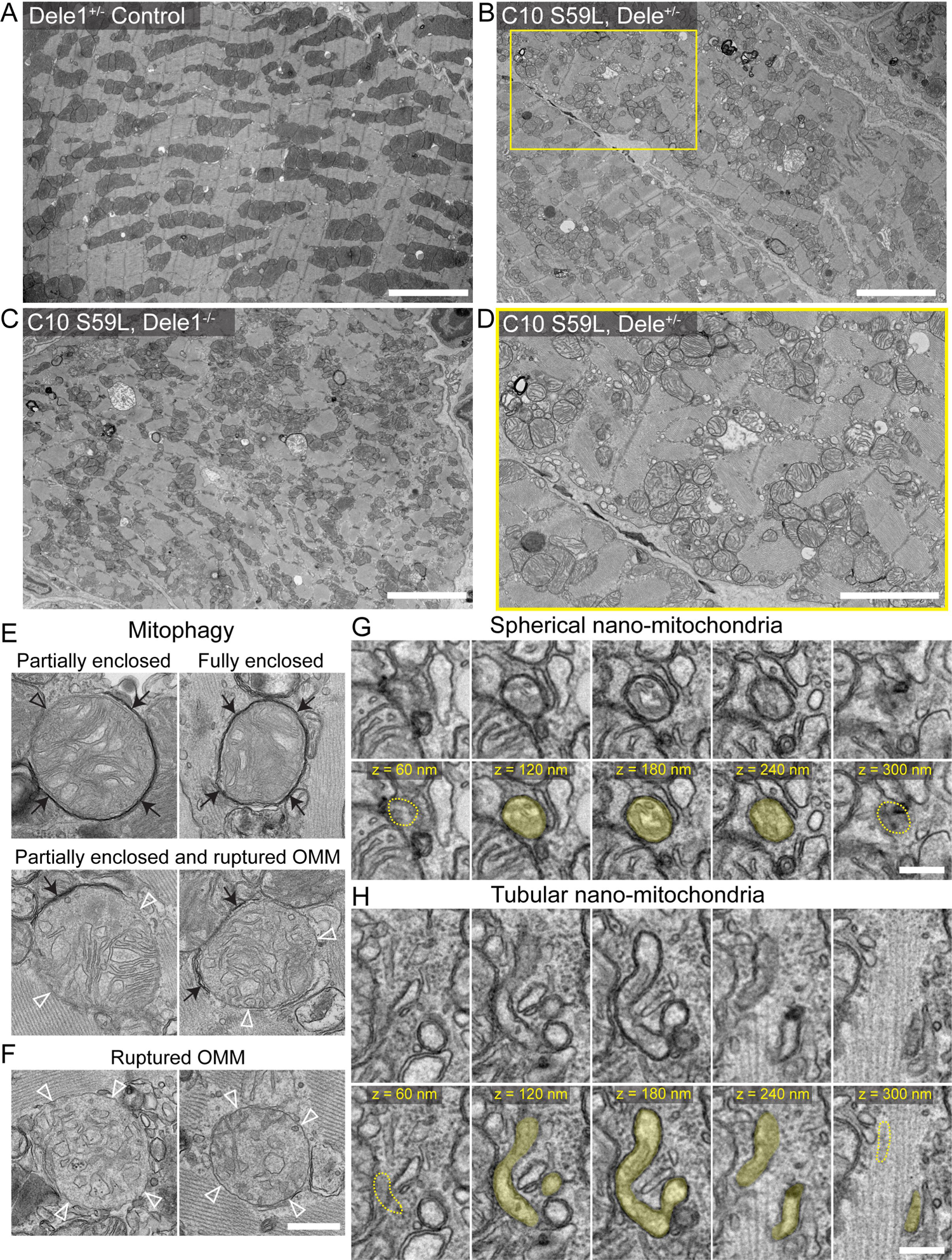
TEM of myocardium and ultrastructural features of mitochondria in C10 S59L on Dele1+/- and Dele1 KO backgrounds. (A-C) Representative TEM images acquired at 2000x direct magnification show areas of myocardium of indicated genotype used for analysis of mitochondria. Scale bar = 5 µm. (D) Image of the subarea boxed yellow in B, acquired at 5000x direct magnification and representative of images used to quantify ultrastructural features of mitochondria detailed in Figure 4. Scale bar = 2.5 µm. (E) Examples of mitochondria that are partially or fully enclosed by electron-dense phagosome membranes (black arrows). Open black arrowhead indicates a portion of the mitochondria that is not enclosed. Partially enclosed mitochondria with ruptured OMMs were also observed. Open white arrowheads indicate sites where the intact IMM is visible, but the OMM is absent. (F) Examples of mitochondria with ruptured OMMs. Open white arrowheads indicate sites where the intact IMM is visible, but an OMM is absent. Scale bar in F = 500 nm and applies to E. (G) Serial sections through a 250 nm diameter mitochondrion show that it is a spherical nano-mitochondrion spanning fewer than five 60-nm sections (< 300 nm in Z). Top row shows the five serial sections without colorization, bottom row shows the same serial sections with the nano-mitochondrion shaded yellow. Yellow dotted lines indicate absence of the mitochondrion in neighboring serial sections. Scale bar = 200 nm. (H) Five serial sections of 60-nm thickness show a 100 nm-wide tubule-shaped mitochondrion. Top row shows five serial sections through the tubular nano-mitochondrion, bottom row shows the same serial sections with the tubular nano-mitochondrion shaded yellow. The yellow dotted lines indicate absence of the mitochondrion in the neighboring section. Scale bar = 200 nm.

**Supplemental Figure 7.**
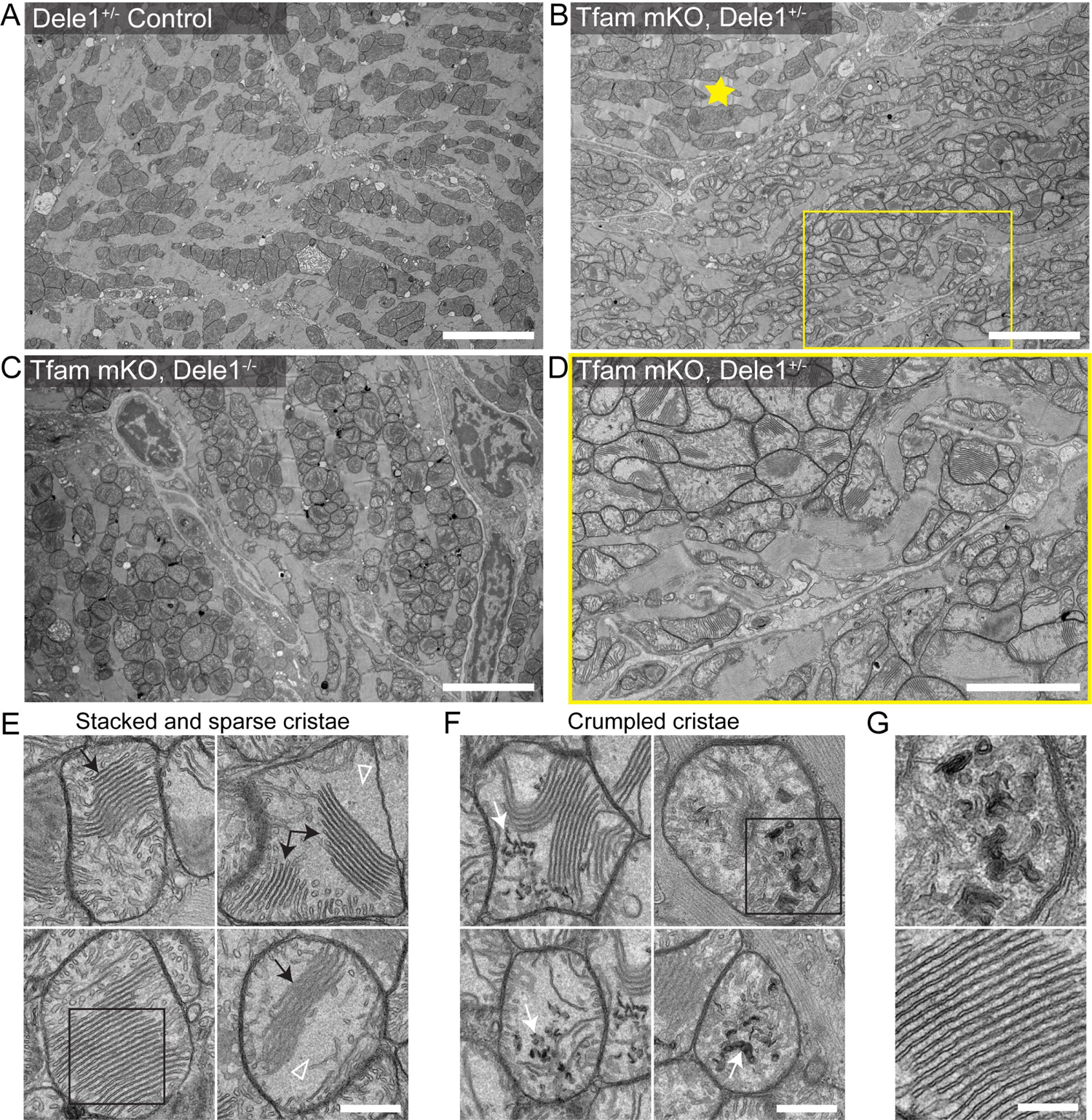
TEM of myocardium and ultrastructural features of mitochondria in *Tfam* mKO on *Dele1*+/- and *Dele1* KO backgrounds. (A-C) Representative TEM images acquired at 2000x direct magnification show areas of myocardium of indicated genotype used for analysis of mitochondria. Yellow star in B indicates a myocyte with milder structural phenotype compared to neighboring myocytes, illustrating the observed mosaicism of the phenotype. Scale bar = 5 µm. (D) Image of the subarea boxed yellow in B, acquired at 5000x direct magnification and representative of images used to quantify ultrastructural features of mitochondria detailed in Figure 4. Scale bar = 2.5 µm. (E) *Tfam* mKO mitochondria displayed populations of closely aligned “stacked” cristae (black arrows) and sparse areas filled with a granular matrix material and few cristae (open white arrowheads). Scale bar = 500 nm. (F) Examples of crumpled cristae (white arrows) that occurred in *Tfam* mKO mitochondria. Scale bar = 500 nm. (G) Stacked cristae boxed in E and crumpled cristae boxed in F are shown enlarged in G. Scale bar = 250 nm.

**Supplemental Figure 8.**
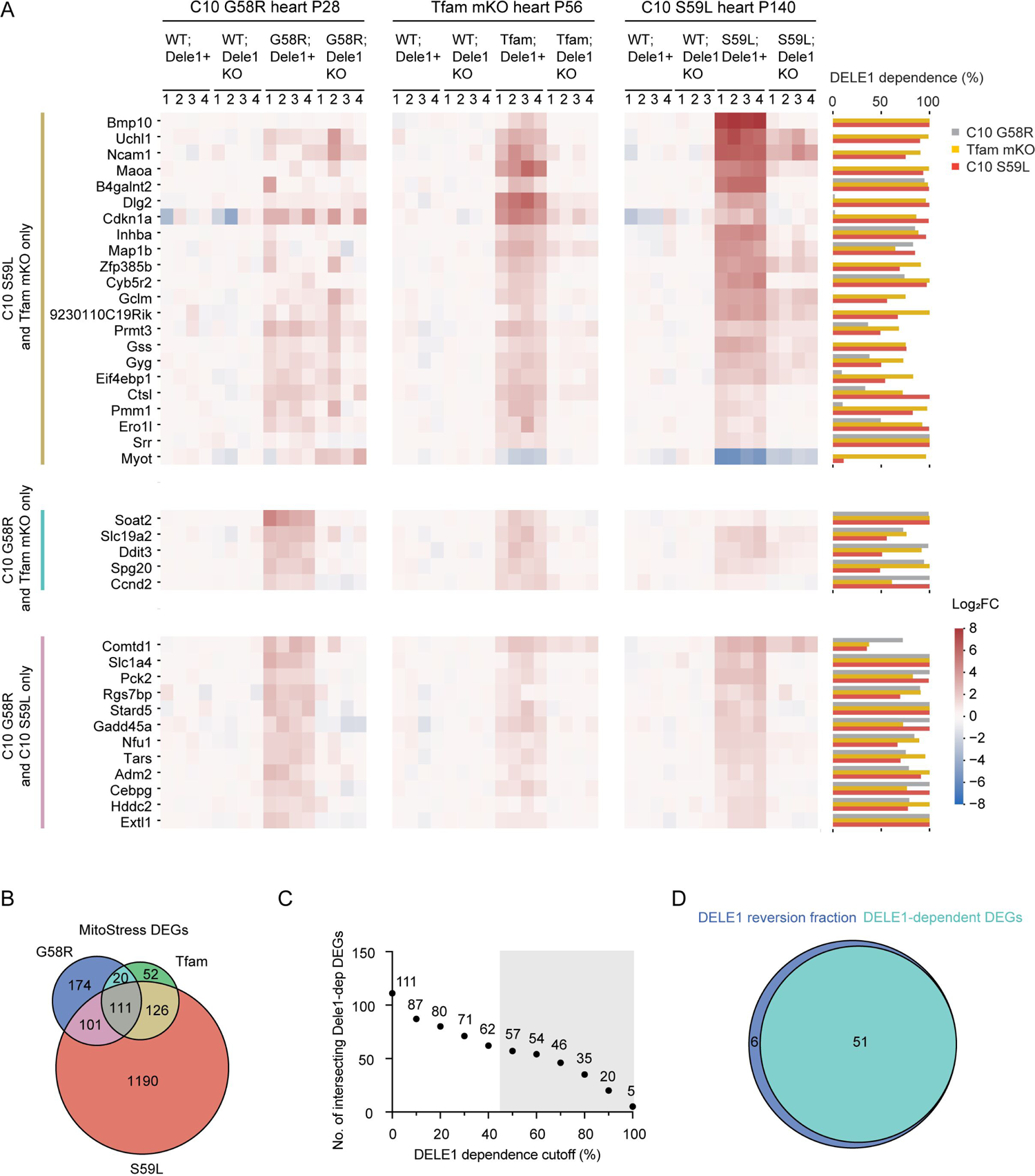
DELE1 mt-ISR transcriptional response in heart is similar in response to diverse mitochondrial stressors. (A) Heat map of Log_2_FC for DELE1-dependent DEGs detected in hearts of two out of three myopathy/cardiomyopathy models. (B) Venn diagram showing intersection of stress-induced DEGs in heart among the 3 models of myopathy/cardiomyopathy. (C) Plot depicts the number of stress induced DEGs ≥ indicated cutoffs for percent DELE1 dependence. (D) Venn diagram showing intersection of DELE1-dependent DEGs and mitochondrial stress-induced DEGs that are ≥ 50% DELE1-dependent.

**Supplemental Figure 9.**
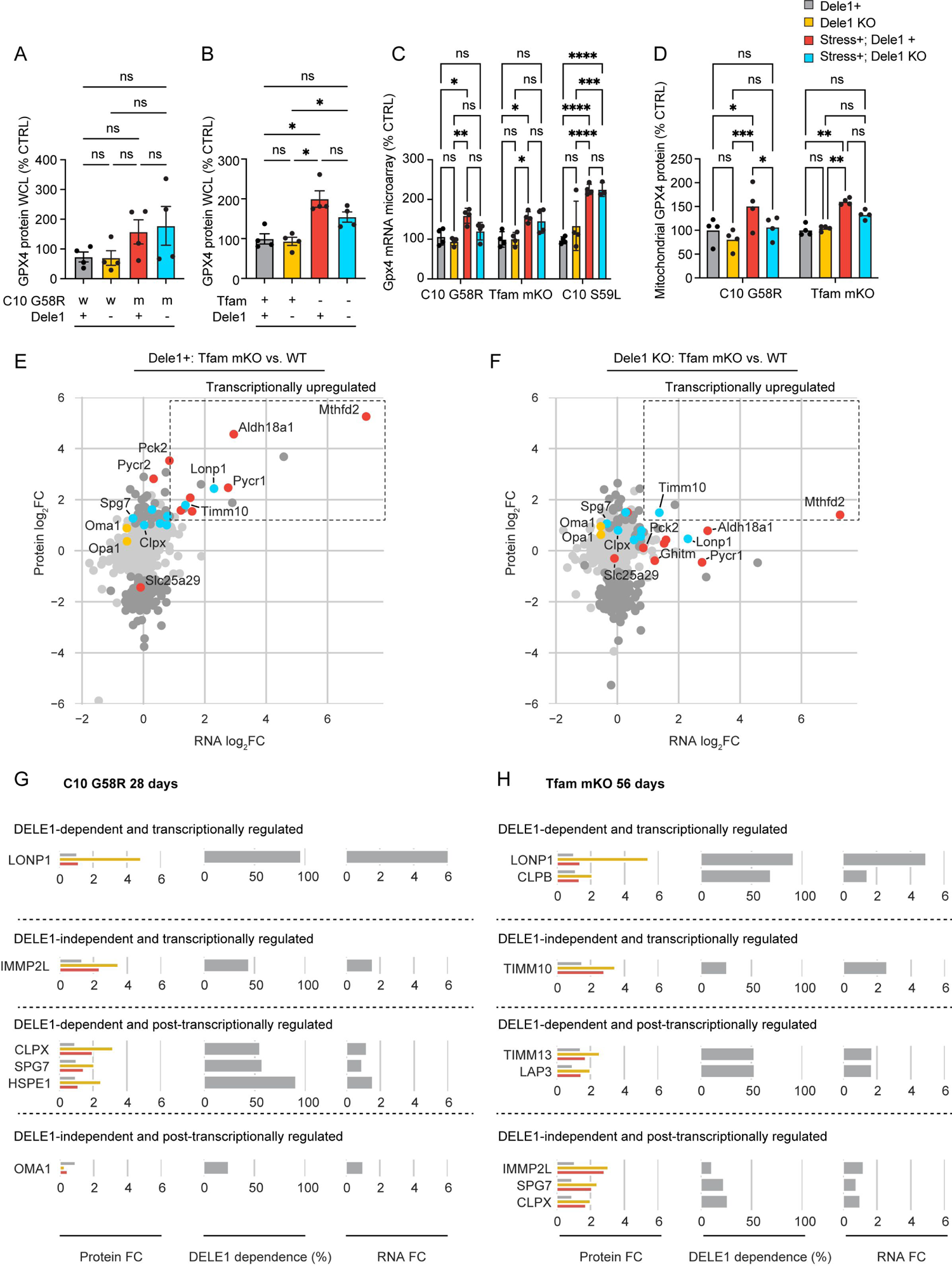
DELE1 mt-ISR transcriptional response in heart has variable effects on *Gpx4* expression and mediates most transcriptionally driven increases in mitochondrial proteins involved in proteostasis. (A-B) GPX4 protein levels from heart whole cell lysate (WCL) of indicated genotypes. (C) *Gpx4* mRNA levels measured in microarray experiments, described in (Fig 5C). (D) GPX4 mitochondrial protein levels measured in proteomics experiments, described in (Fig 5G and H). (E - F) Scatterplot compares RNA log_2_FC for *Tfam* mKO vs. control (in the presence of DELE1) and mitochondrial protein log_2_FC for *Tfam* mKO vs. control animals in the presence of DELE1 (left) or the absence of DELE1 (right). (G – H) Bar graphs showing protein and RNA fold changes for proteins annotated as proteases and chaperones in MitoCarta3.0, from experiments described in (Fig. 5C, G, and H).

**Supplemental Figure 10.**
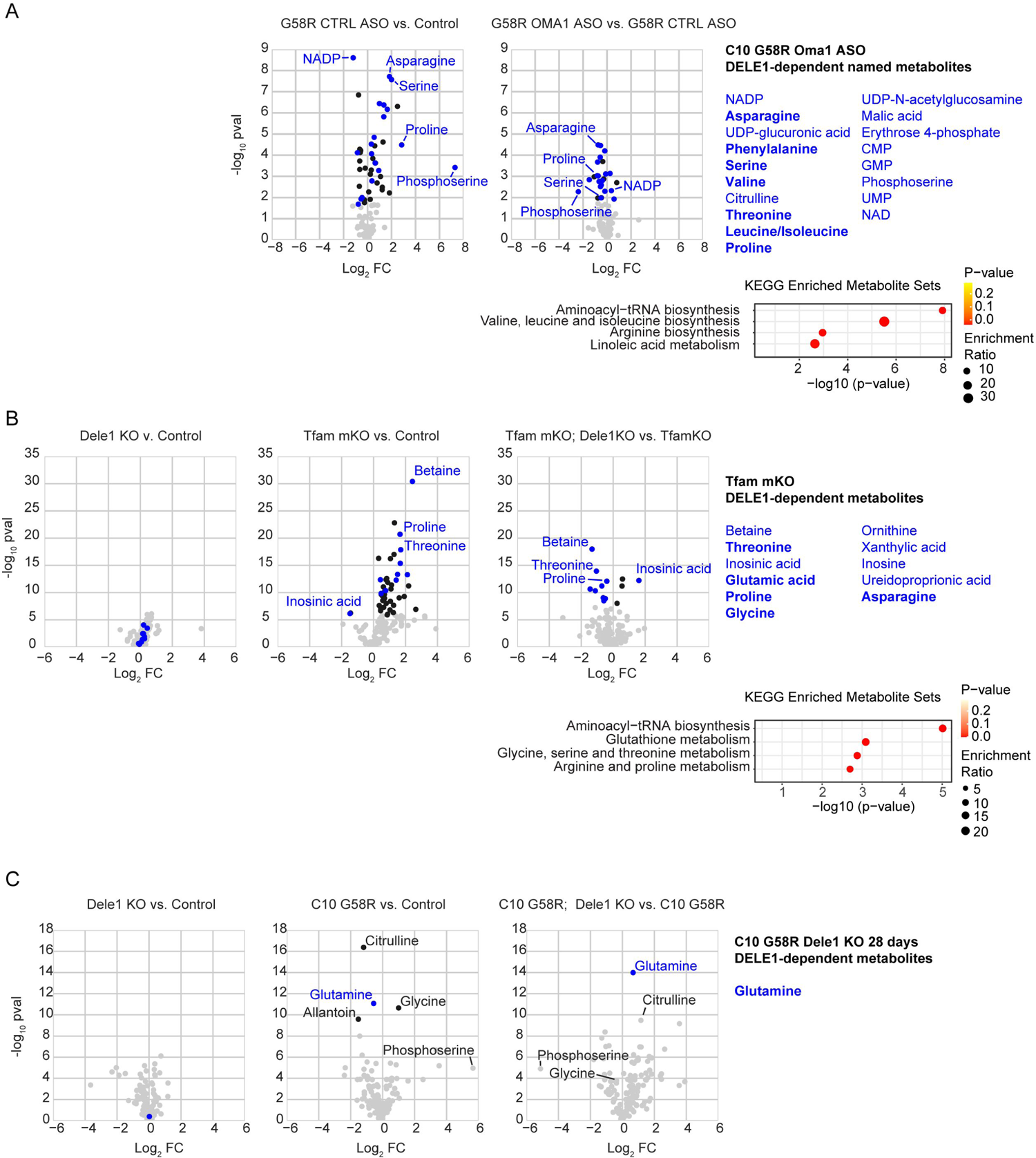
Metabolomics from hearts of C10 G58R and *Tfam* mKO models of mitochondrial myopathy/cardiomyopathy. (A - C) Volcano plots of metabolites identified in targeted (B and C) or untargeted (A) metabolomics experiments and enrichment analysis among KEGG metabolite sites for the DELE1-dependent metabolites. Only named features in the untargeted metabolomics data are plotted in (A). DELE1-dependent metabolites are in blue, with amino acids bolded.

**Supplemental Figure 11.**
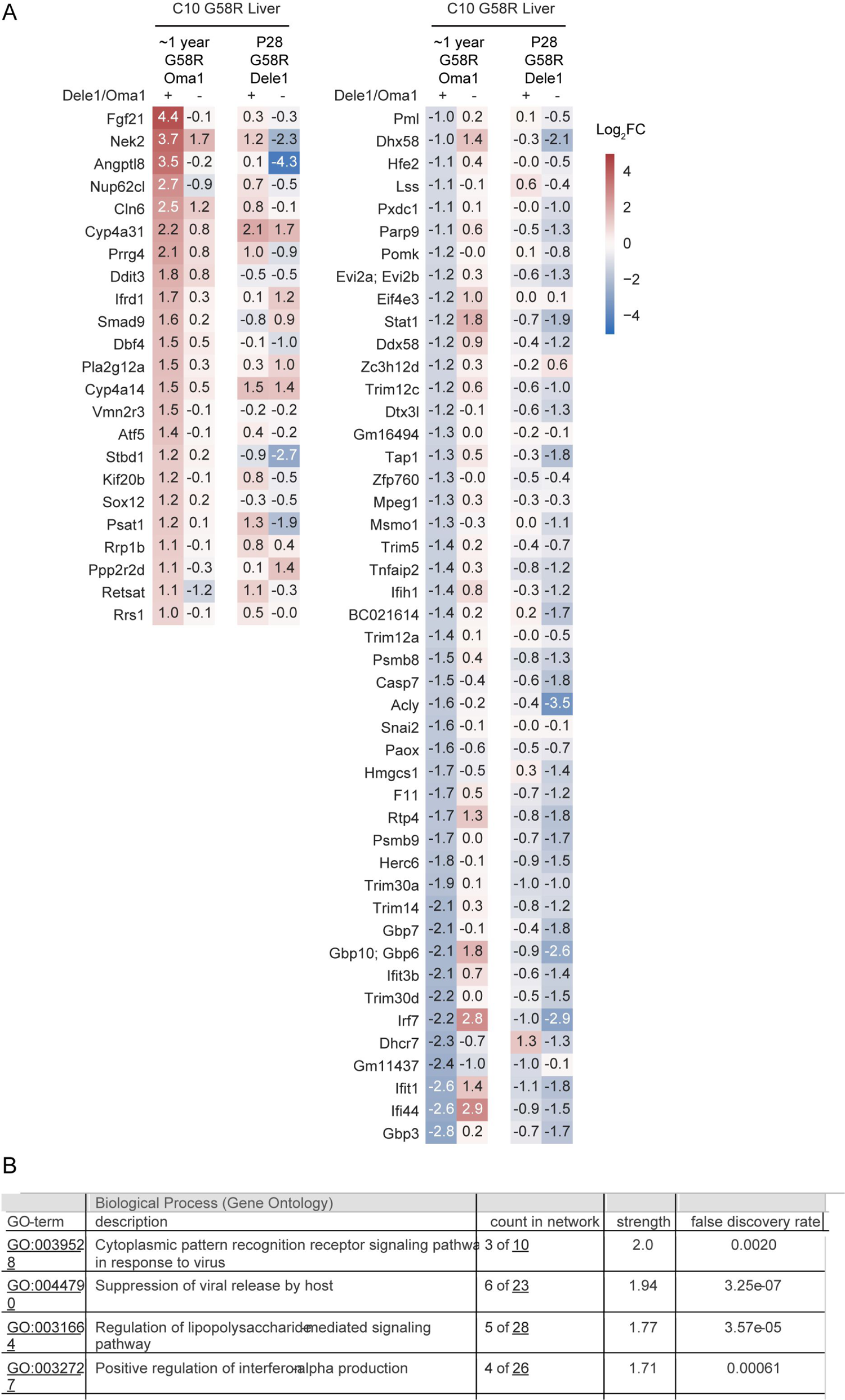
Transcriptomics from liver from C10 G58R mice with either *Dele1* KO (P28) or *Oma1* knockdown using an ASO (∼1 year). (A) Heat map depicts the OMA1-dependent DEGs detected in liver from ∼1 year old C10 G58R mice compared to P28 C10 G58R mice. No DELE1-dependent DEGs were detected at P28. (B) Top gene ontology terms for the OMA1-dependent DEGs from livers of ∼1 year old C10 G58R mice in (A).

**Supplemental Figure 12.**
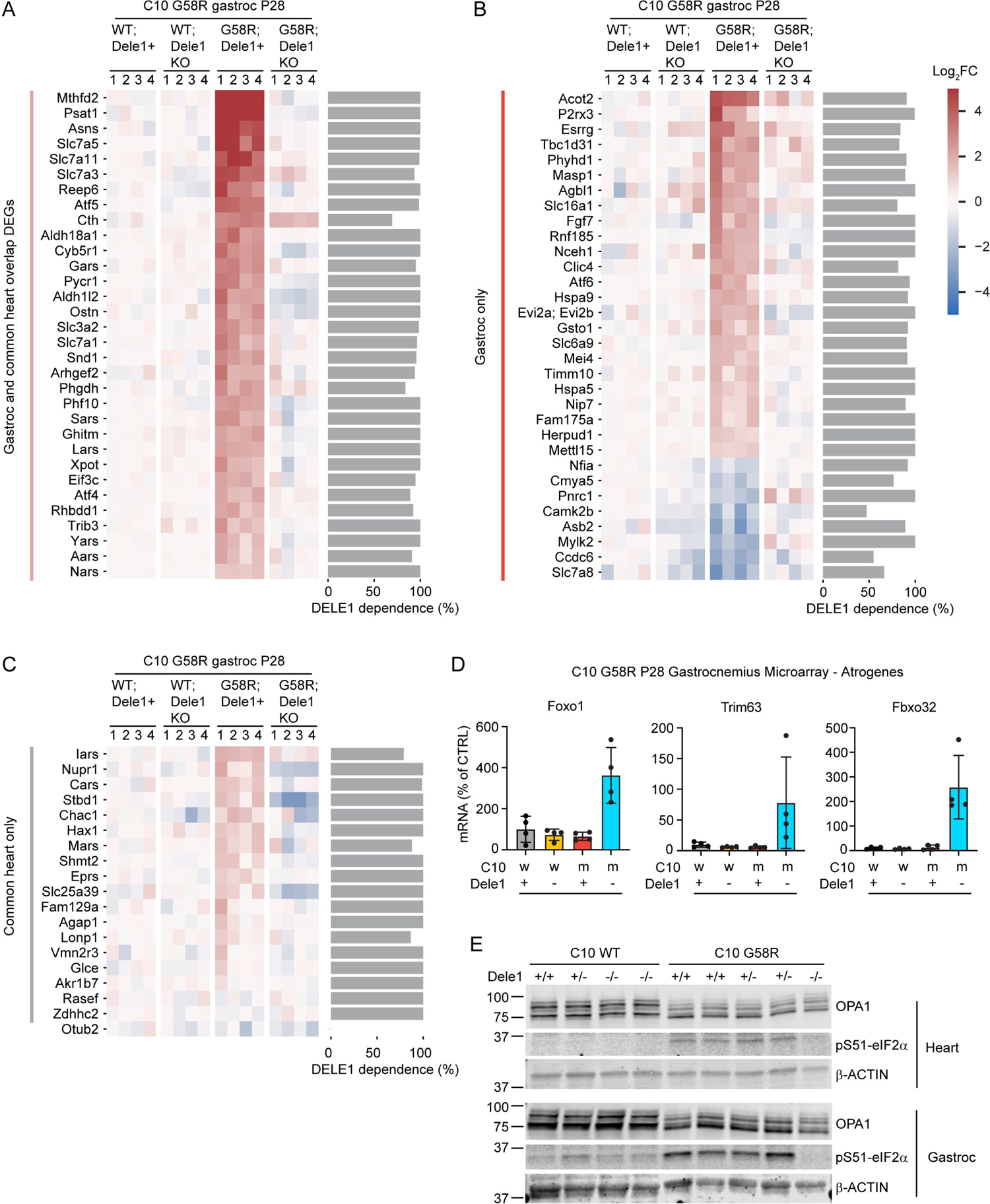
Transcriptomics from gastrocnemius skeletal muscle from P28 C10 G58R; *Dele1* KO mice and littermates. (A) Heatmap of gene expression changes from gastrocnemius skeletal muscle for intersection of DELE1-dependent DEGs in gastrocnemius skeletal muscle and heart DELE1 mt-ISR signature. Bargraph on right represents percent DELE1 dependence. (B) Heatmap of gene expression changes from gastrocnemius skeletal muscle for DELE1-dependent DEGs in gastrocnemius skeletal muscle that are not part of heart DELE1 mt-ISR signature. Bargraph on right represents percent DELE1 dependence. (C) Heatmap of gene expression changes from gastrocnemius skeletal muscle for genes in the heart DELE1 mt-ISR signature that were not significantly DELE1-dependent in the gastrocnemius muscle. Bargraph on right represents percent DELE1 dependence. (D) Bar graphs depicting mRNA changes for *Foxo1* and two canonical atrogenes, *Trim63* and *Fbox32*, measured in microarray data of gastrocnemius skeletal muscle lysates. (E) Immunoblot of heart (top) and gastrocnemius muscle (bottom) from P28 C10 G58R; *Dele1* KO mice and their littermates, showing DELE1-dependent phosphorylation of eIF2α.

**Supplemental Figure 13.**
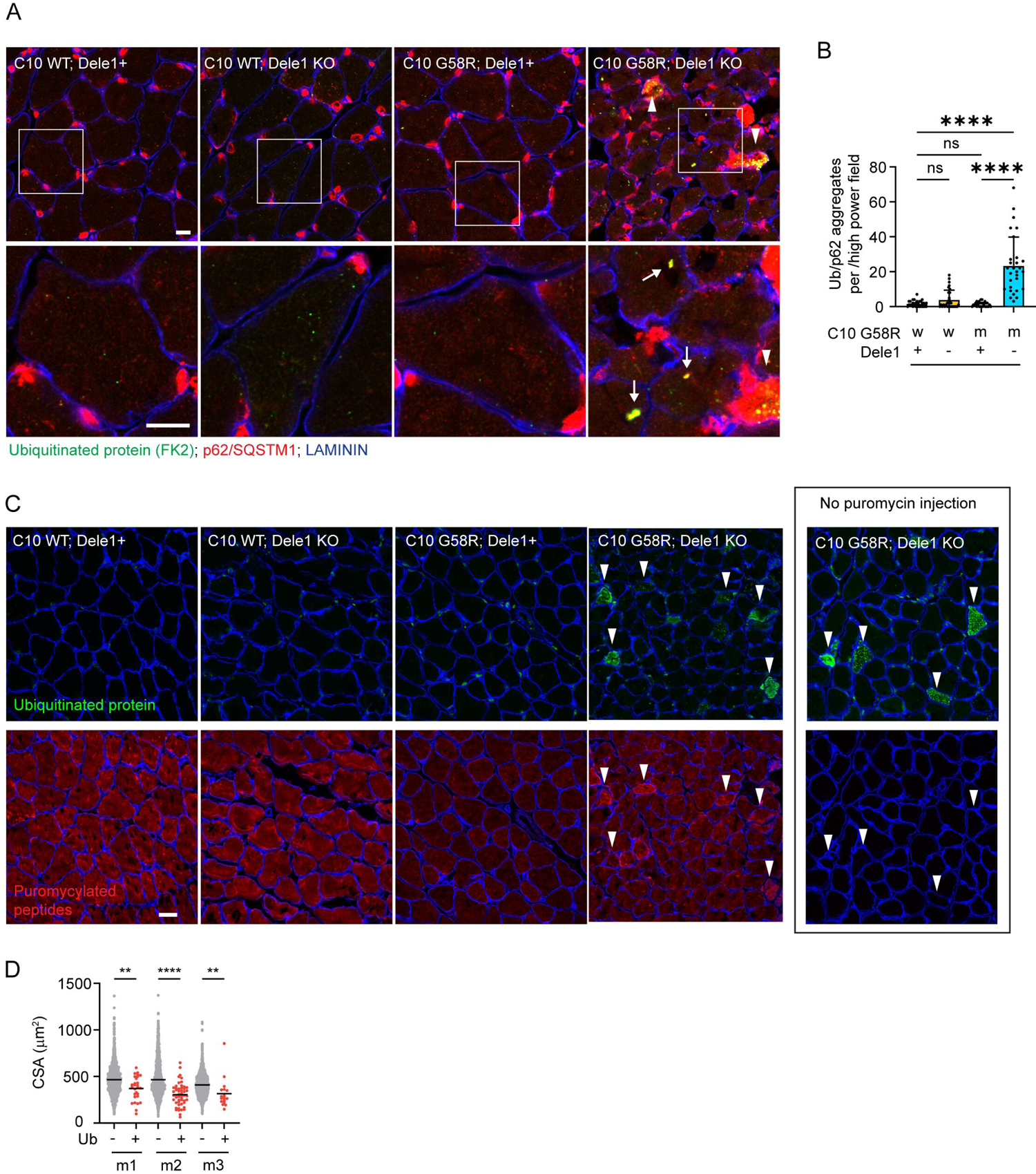
The Dele1 mt-ISR prevents disruptions in translation-associated proteostasis. (A) Representative immunofluorescence images of gastrocnemius muscle from C10 G58R; *Dele1* KO mice and littermates, showing fibers with confluent aggregates of ubiquitinated proteins co-localized with the aggregate-forming adaptor protein p62, suggesting proteostatic collapse (arrow heads) and individual aggregates staining positive for ubiquitinated protein and the aggregate-forming adaptor protein p62 (arrows) in high power (60X) images (bottom panels). Scale bars = 10 μm. Note: animals were not injected with puromycin in this experiment. (B) Quantification of (B). Aggregates positive for both p62 and ubiquitinated protein immunofluorescence were counted in 10 high power (60X) fields. N = 3 mice per genotype with 29 or 30 fields counted total per sample. High-power (60X) field size is 132.58 μm X 132.58 μm. (C) Representative immunofluorescence images of gastrocnemius muscle from P28 C10 G58R; Dele1 KO mice and littermates injected with puromycin 30 min prior to sacrifice as in (Fig. 7E). Muscle cross-sections were immunostained for ubiquitinated proteins (using the FK2 antibody) (green), puromycin (red), and LAMININ (blue). Arrow heads indicate muscle fibers containing many or confluent aggregates of ubiquitinated protein that were also co-stained for elevated puromycylated polypeptides. N = 1 mouse for each genotype except for P28 C10 G58R; Dele1 KO mice for which N = 3 mice. Scale bars = 20 μm. (D) Quantification of myofiber cross-sectional area (CSA) in (Fig. 7E). The average CSA for Ub+ and Ub-myofiber is shown in graph separately for three mice (m1 – 3) in graph. N = 3 mice with 10 high power fields counted per mouse.

### Supplemental Tables

**Table 1.** Transcriptomics from hearts of three myopathy/cardiomyopathy mouse model.

**Table 2.** Mitochondrial proteomics from hearts of three myopathy/cardiomyopathy mouse model.

**Table 3.** Metabolomics from hearts of two myopathy/cardiomyopathy mouse model, *Tfam* mKO and C10 G58R.

